# Tracking the Ancestry of Known and ‘Ghost’ Homeologous Subgenomes in Model Grass *Brachypodium* Polyploids

**DOI:** 10.1101/2021.05.31.446465

**Authors:** Rubén Sancho, Luis A. Inda, Antonio Díaz-Pérez, David L. Des Marais, Sean Gordon, John Vogel, Joanna Lusinska, Robert Hasterok, Bruno Contreras-Moreira, Pilar Catalán

**Affiliations:** Department of Agricultural and Environmental Sciences, High Polytechnic School of Huesca, University of Zaragoza, Huesca, Spain; Grupo de Bioquímica, Biofísica y Biología Computacional (BIFI, UNIZAR), Unidad Asociada al CSIC, Spain; Instituto Agroalimentario de Aragón (IA2), Universidad de Zaragoza, Spain; Instituto de Genética, Facultad de Agronomía, Universidad Central de Venezuela, Venezuela; The Arnold Arboretum of Harvard University, Boston, MA, USA; Department of Civil and Environmental Engineering, Massachusetts Institute of Technology, Cambridge, MA, USA; DOE Joint Genome Institute, Berkeley, CA, USA; The University of California at Berkeley, Berkeley, CA, USA; Plant Cytogenetics and Molecular Biology Group, Institute of Biology, Biotechnology and Environmental Protection, Faculty of Natural Sciences, University of Silesia in Katowice, Katowice, Poland; Department of Genetics and Plant Breeding, Estación Experimental de Aula Dei-Consejo Superior de Investigaciones Científicas, Zaragoza, Spain; Ensembl Plants, European Bioinformatics Institute, EMBL-EBI, Hinxton, UK; Tomsk State University, Tomsk, Russia

**Keywords:** allopolyploids, autopolyploids, chromosomal barcodes, ‘ghost’ progenitor genomes, phylogenomic subgenome detection pipeline

## Abstract

Unraveling the evolution of plant polyploids is a challenge when their diploid progenitor species are extinct or unknown or when their progenitor genome sequences are unavailable. The subgenome identification methods cannot adequately retrieve the homeologous genomes that are present in the allopolyploids if they do not take into account the potential existence of unknown progenitors. We addressed this challenge in the widely distributed dysploid grass genus *Brachypodium*, which is a model genus for temperate cereals and biofuel grasses. We used a transcriptome-based phylogeny and newly designed subgenome detection algorithms coupled with a comparative chromosome barcoding analysis. Our phylogenomic subgenome detection pipeline was validated in *Triticum* allopolyploids, which have known progenitor genomes, and was used to infer the identities of three extant and four ‘ghost’ subgenomes in six *Brachypodium* polyploids (*B. mexicanum, B. boissieri, B. retusum, B. phoenicoides, B. rupestre and B. hybridum*), of which five contain undescribed homeologous subgenomes. The existence of the seven *Brachypodium* progenitor genomes in the polyploids was confirmed by their karyotypic barcode profiles. Our results demonstrate that our subgenome detection method is able to uncover the ancestral genomic components of both allo- and autopolyploids.

## Introduction

While the genomic origins of some polyploid plants have been deduced using comparative genomics (e.g. wheats (Marcussen et al. 2014a; Appels et al. 2018), deciphering the genomic history of many allopolyploids has proven to be challenging when the progenitor species are extinct or unknown (Soltis and Soltis 2016) or when the contributing parental genomes are highly similar (Brassac and Blattner 2015; Kamneva et al. 2017). Incomplete genome assemblies further complicate the delineation of homeologous genomes in allopolyploid plants, which is a typical scenario in angiosperms except for a few experimental plants and crops (Soltis et al. 2016; Scholthof et al. 2018). Multispecies coalescent species trees and networks, together with syntenic read-mapping phylogenetic approaches, have successfully reconstructed the history of the homeologous genomes of some allopolyploid plants (Bombarely et al. 2014; Bertrand et al. 2015; Marcussen et al. 2015; Novikova et al. 2016; Oxelman et al. 2017). However, most of the studied cases correspond to allopolyploids with known diploid genome donors. Few studies have identified subgenomes that were derived from unknown diploid ancestors (Kamneva et al. 2017) or have explicitly incorporated both known and unknown ‘ghost’ subgenomes into the searching strategy (Marcussen et al. 2015). Coalescence-based methods account for incomplete lineage sorting (ILS) events across the gene trees and retrieve consensus merging scenarios for the subgenomes for each allopolyploid (Marcussen et al. 2015; Kamneva et al. 2017). Nonetheless, these protocols are challenging due to the computational overhead for the likelihood or Bayesian-based methods or are only currently available for allotetraploids (e.g., AlloppNET and AlloppMUL models; Jones 2017). Additionally, selecting the optimal hybridization scenarios is impeded in cases in which the progenitor diploid genomes are unknown and the number of possible subgenome combinations increases with ploidy (Bertrand et al. 2015; Marcussen et al. 2015).

Allopolyploids are common in the grass family and account for 70% of the current species (Stebbins 1949; Kellogg 2015). *Brachypodium* was selected as a model system for cereals and biofuel grasses (Scholthof et al. 2018; Catalán and Vogel 2020). This pooid genus has become an indispensable tool for investigating many aspects of the functional genomics, biology and evolution of grasses and monocots more broadly and translating the fundamental biological insights to crop species (Catalán and Vogel 2020). Recent phylogenetic studies have suggested that allopolyploidy has been a prevalent speciation mechanism in *Brachypodium* (Catalán et al. 2016; Díaz-Pérez et al. 2018) and, indeed, that most allopolyploid *Brachypodium* species likely resulted from crosses of dysploid progenitor species that had different basic chromosome numbers (Betekhtin et al. 2014; Díaz-Pérez et al. 2018). The best-known case is the annual allotetraploid *B. hybridum* (2n=30, x=10+5), which was derived from the cross and subsequent genome doubling of the diploid *B. stacei-type* (2n=20, x=10) and *B. distachyon-type* (2n=10, x=5) progenitors (Catalán et al. 2012; López-Álvarez et al. 2012; Catalán et al. 2014; Shiposha et al. 2019; Gordon et al. 2020). The re-creation of a stable synthetic allotetraploid that phenotypically resembles the natural *B. hybridum* corroborated the allopolyploid origin of this neopolyploid species (Dinh Thi et al. 2016). In contrast, the evolutionary history of the perennial *Brachypodium* allopolyploids is more intriguing due to their full or partial ‘ghost’ homeologous subgenomes, which have only been studied with a limited set of nuclear and plastid loci (Catalán et al. 2016; Díaz-Pérez et al. 2018).

Here, we present a novel approach (PhyloSD) for uncovering the homeologous subgenomes that are present in the *Brachypodium* allopolyploids and reconstruct their evolution focusing specifically on species whose diploid progenitors are extinct or unknown (‘ghost’ subgenomes). We used the well-known phylogeny and diploid progenitor genomes of the allopolyploid *Triticum* species (Marcussen et al. 2014a) to benchmark our algorithms. Then, we applied the algorithms to *Brachypodium* in an attempt to retrieve its reticulate history by focusing on six putative polyploids. We used our subgenome detection algorithms as an *a priori* assignment of homeologs to the genomes of their hypothetical diploid progenitors. We further validated the computational pipeline and reconstructed a robust phylogeny for the genomes and homeologous subgenomes of 12 *Brachypodium* species and ecotypes. These computational approaches were validated using fluorescence *in situ* hybridization (FISH)-based comparative chromosome barcoding (CCB), which enables the specific painting of whole chromosomes or their regions. The CCB proved to be effective in tracking the structural and evolutionary trajectories of individual chromosomes and whole karyotypes in some dicots, (e.g., *Arabidopsis thaliana* and its relatives; Lysak et al. 2006) and monocots (e.g., rice; Hou et al. 2018) and recently contributed to dissecting the karyotype organization of some *Brachypodium* species (Lusinska et al. 2019). Here, this combined phylogenomic and cytomolecular strategy enabled us to propose hypotheses about the identities of the known and unobserved progenitor genome donors in the studied *Brachypodium* polyploid species and to infer their times of origin.

## Results

### The Phylogenomic Subgenome Detection (PhyloSD) pipeline

PhyloSD employs three sequential algorithms: *Nearest Diploid Species Node, Bootstrapping Refinement and Subgenome Assignment* (fig. 1A-C). The input is a set of pre-computed multiple sequence alignments (MSAs) of coding sequences and transcripts that contain the ingroup diploid orthologs and polyploid homeologs and outgroup orthologs. The pipeline consists of (i) a computational filtering step, (ii) labeling the homeologs, and (iii) allocating the homeologs to subgenomes (Supplementary Material; supplementary fig. S1).

**Fig. 1.**
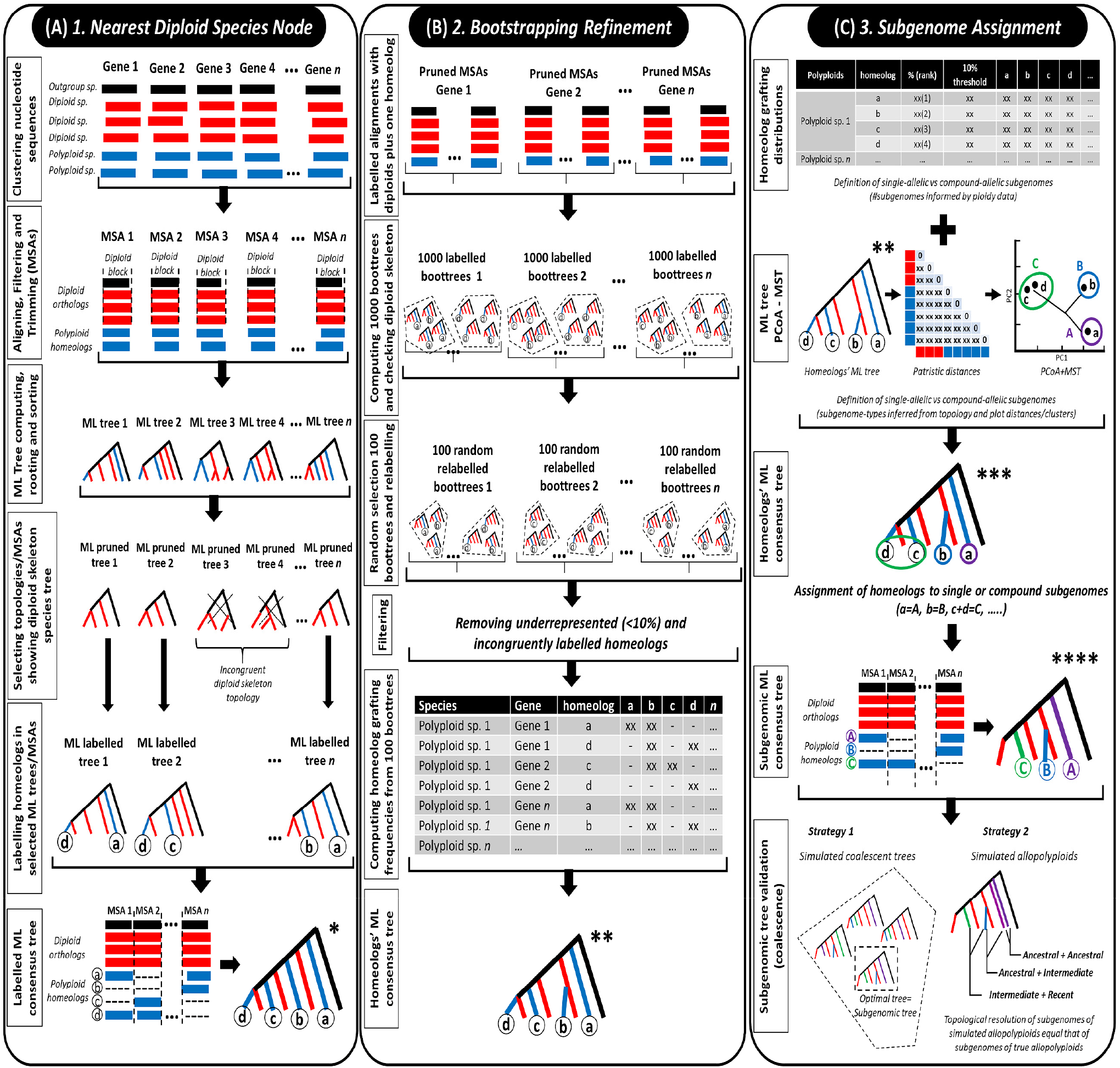
A summarized workflow of the Phylogenetic Subgenome Detection (PhyloSD) pipeline highlighting the three *Nearest Diploid Species Node* **(A)**, *Bootstrapping Refinement* **(B)** and *Subgenome Assignment* **(C)** algorithms. Black, red and blue colors indicate the diploid outgroup and diploid and polyploid ingroup sequences, respectively. The Labeled ML consensus tree (*) that was obtained from the *Nearest Diploid Node* algorithm (a) was fine-tuned by the *Bootstrapping Refinement* algorithm (b) resulting in the Homeologs’ ML consensus tree (**). The Homeologs’ ML consensus tree (**) was readjusted by the *Subgenome Assignment* algorithm (c) (***) and genomically relabeled resulting in the Subgenomic ML consensus tree (****). The MSA (Multiple Sequence Alignment); ML (Maximum-Likelihood); PCoA-MST (Principal Coordinate Analysis and superimposed Minimum Spanning Tree).

The *Nearest Diploid Species Node* algorithm labels the homeologous sequences according to their grafting positions with respect to the nearest diploid species in the optimal diploid skeleton tree and its stem branch (fig. 1A). MSAs with missing diploid sequences and non-overlapping alignment blocks are discarded. Maximum Likelihood (ML) phylogenetic trees are subsequently estimated for each of the curated MSAs, thereby obtaining exploratory gene trees (fig. 1A). These trees are further filtered, keeping only the most frequent partitions that have a diploid skeleton topology that is congruent with that of the diploid species tree. The diploid species tree was obtained from parallel coalescent analyses (ASTRAL, STEAC, STAR). Then, the homeologs are labeled in each partition tree according to their grafting positions with respect to the nearest diploid species in the optimal diploid skeleton tree using *ad-hoc* labeling rules (‘a’ to ‘i’; figs. 1A, 2A, 3A) and assuming that each homeolog type would represent a subgenome in the polyploid. A labeled ML consensus tree is then computed from all of the labeled partitions (fig. 1A; supplementary fig. S1).

**Fig. 2.**
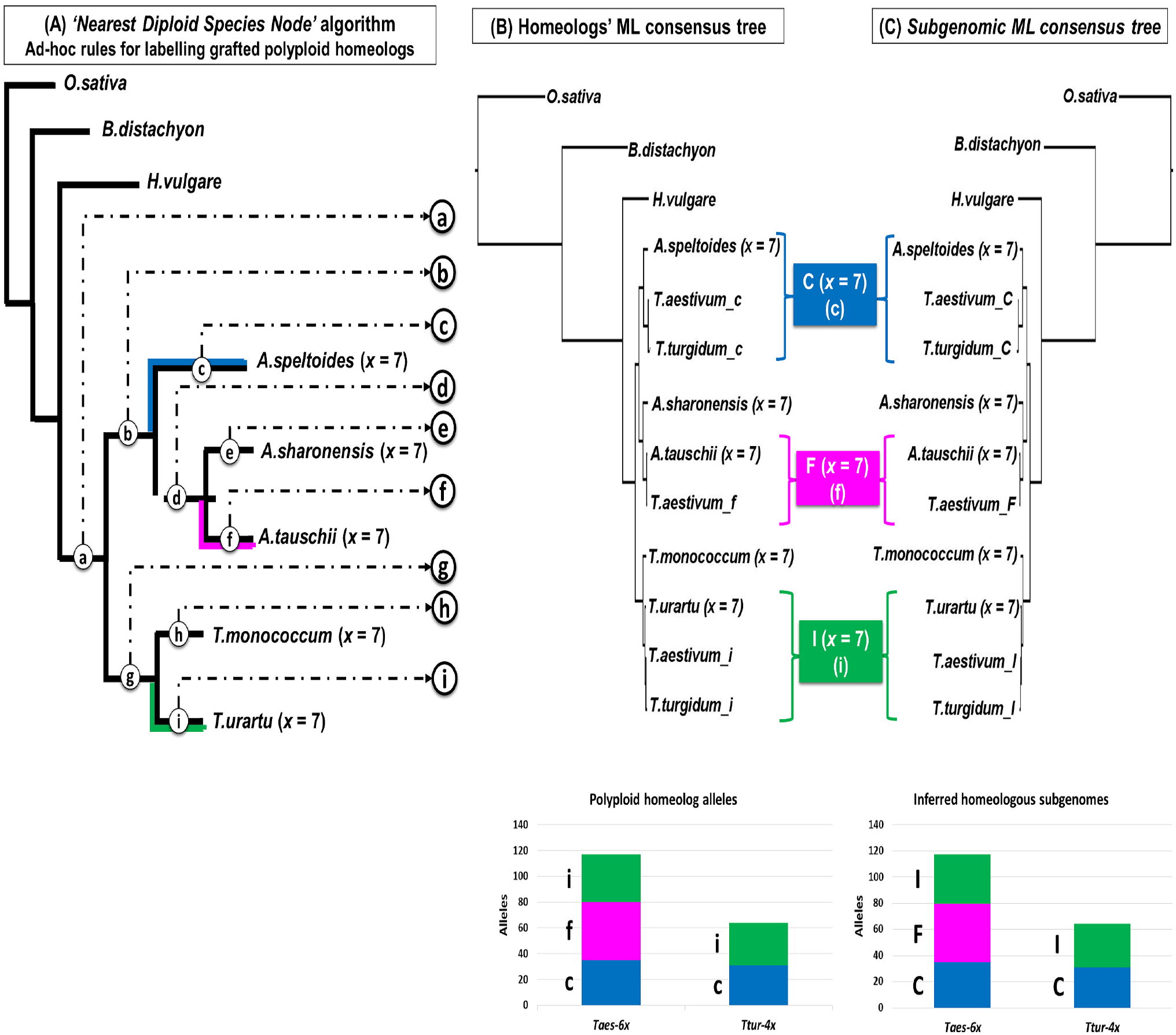
**(A)** Schematic *Triticum-Aegilops* tree illustrating the diploid skeleton tree (thick black branches) of the orthologous diploid genome sequences (*A. speltoides, A. sharonensis, A. tauschii, T. monococcum, T. urartu*) of x=7 showing the *ad hoc* labeling rules (lowercase letters, ‘a’-‘i’) for the grafting positions of the *Triticum* polyploid homeolog sequences according to the *Nearest Diploid Species Node* algorithm; **(B)** *Triticum-Aegilops* homeologs’ ML consensus tree based on 48 core genes and 181 homeologs (table 1A) with the polyploid homeolog sequences labeled according to the *Nearest Diploid Species Node* algorithm (‘c’, ‘i’, ‘f’); **(C)** *Triticum-Aegilops* subgenomic ML consensus tree based on 48 core genes with the homeolog subgenomes labeled according to the *Subgenome Assignment* algorithm (‘C’, ‘F’, ‘I’) (table 1A; supplementary fig. S3). *Oryza sativa, Brachypodium distachyon* and *Hordeum vulgare* were used as the outgroups. Asterisks indicate branches with SH-aLRT/UltraFast Bootstrap support (BS) <80/95; the remaining branches have 100/100 values. The bar diagrams represent the frequencies of the homeologs in each polyploid and their assignments to the homeologous subgenomes (table 1A).

**Fig. 3.**
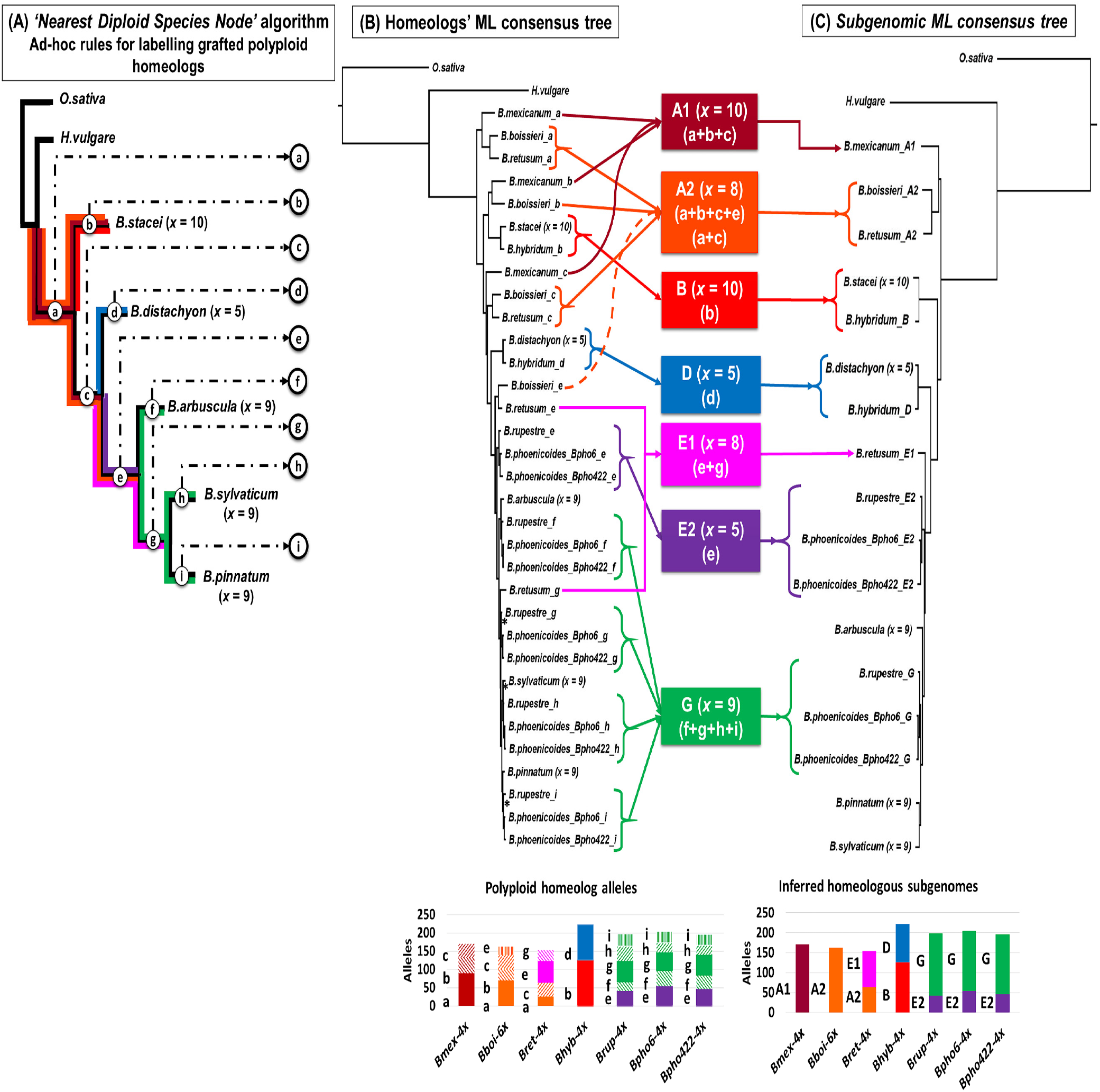
**(A)** Schematic *Brachypodium* tree illustrating the diploid skeleton tree (thick black branches) of the orthologous diploid genome sequences and their respective chromosome base numbers (*B. stacei* x=10; *B. distachyon* x=5; core perennial *B. arbuscula, B. sylvaticum* and *B. pinnatum* clade x=9) and the nesting positions of the *Brachypodium* polyploid homeolog sequences showing the *ad hoc* labeling rules (lowercase letters, ‘a’-‘i’) for the grafting positions of the *Brachypodium* polyploid homeolog sequences according to the *Nearest Diploid Species Node* algorithm; **(B)** *Brachypodium* homeologs’ ML consensus tree based on 322 core transcripts and 1,307 homeologs (table 1B) with the polyploid homeolog sequences labeled according to the *Nearest Diploid Species Node* algorithm (‘a’, ‘b’, ‘c’, ‘d’, ‘e’, ‘f’, ‘g’, ‘h’, ‘i’); **(C)** *Brachypodium* subgenomic ML consensus tree based on 322 core genes with the homeolog subgenomes labeled according to the *Subgenome Assignment* algorithm (‘A1’, ‘A2’, ‘B’, ‘D’, ‘E1’, ‘E2’, ‘G’) (table 1B; supplementary fig. S5). *Oryza sativa* and *Hordeum vulgare* were used as the outgroups. Asterisks indicate branches with SH-aLRT/UltraFast Bootstrap supports (BS) <80/95; the remaining branches have 100/100 values. The bar diagrams represent the frequencies of the homeologs in each polyploid and their assignments to the homeologous subgenomes (table 1B).

The *Bootstrapping Refinement* algorithm tests the labeling of the homeologs. Bootstrap analyses are performed to generate patterns of the branch distribution for each homeolog type, assuming that a single homeolog could have been grafted erroneously in topological-vicinity branches by accident. The labeled MSAs from the previous step are pruned and used to generate new datasets, each of which contains all of the diploid orthologs plus one polyploid homeolog at a time (fig. 1B). Next, one thousand labeled ML bootstrapping trees (boottrees) are computed for each pruned alignment. The robustness of the grafted homeologs is assessed using a low bootstrap support cut-off (BS<10%) in each of the consensus boottrees that has a congruent diploid skeleton topology. Poorly represented homeolog types are removed. Then, 100 boottrees are randomly selected from each group and the homeologs are relabeled. The homeolog grafting frequencies from each of the 100 boottrees are computed and the resulting homeologs’ ML consensus tree is constructed (fig. 1B; supplementary fig. S1).

The *Subgenome Assignment* algorithm allocates the homeologs to the corresponding defined polyploid subgenomes based on i) the frequency ranks of their grafting distributions (fig. 1C) and ii) the clusters of homeologs in a principal coordinate analysis (PCoA) with a superimposed minimum spanning tree (MST) plot (fig. 1C). The PCoA-MST is obtained from the pairwise patristic distances, which are computed from the ML consensus tree that was retrieved in the second step (fig. 1B) (supplementary fig. S1). The grafting distributions of the homeologs are evaluated to determine their circumscription to a single or a few contiguous branches of the species tree (fig. 1C). The homeologs are assigned to single-type (single-allelic) subgenomes if they were grafted to single branches with the highest frequency and the remaining graftings were below the cut-off threshold (≤10% of the main grafting frequency). They are assigned to compound-type (multi-allelic) subgenomes if the secondary and subsequent grafting frequencies are above the threshold. The most frequent homeolog types (‘a’, ‘b’, …) are selected according to the expected number of subgenomes that are suggested by ploidy (two for tetraploids and three for hexaploids) and are re-coded as subgenomes (‘A’, ‘B’,…), and the low frequency homeolog types incompatible with the ploidy level of the polyploid are discarded (figs. 1C, 2B, C, 3B, C). The labeled subgenomic MSAs are used to compute the subgenomic consensus ML tree (fig. 1C).

### Benchmarking the phylogenomic subgenome detection pipeline in the *Triticum-Aegilops* allopolyploid complex

The initial *Triticum-Aegilops* data set consisted of 275 ortholog clusters that were obtained from Marcussen et al. (2014b) and Marcussen et al. (2014a) of which only 48 MSAs with 236 homeologs (having a diploid skeleton topology that was congruent with that of the coalescent-based species tree) remained after the filtering steps of the first algorithm (Supplementary Material; supplementary figs. S2A-C). The homeologs were labeled according to the *ad hoc* rules that are presented in fig. 2A. When the incongruently labeled and underrepresented homeologs were removed from the second algorithm, a total of 48 MSAs and 181 homeologs were obtained. These were used to compute the *Triticum*-*Aegilops* maximum-likelihood (ML) consensus tree (fig. 2B). The grafting distributions of the homeologs (supplementary table S2A) and the PCoA-MST clusterings (supplementary fig. S3) of the third algorithm presented a simple scenario in which all of the selected homeologs were assigned to single subgenomes that contained only one homeolog type (table 1A; supplementary table S2A). Thus, homeolog ‘c’ corresponded to the subgenome C that is present in *T. turgidum* and *T. aestivum*, ‘f’ to the subgenome F that is present in *T. aestivum* and ‘i’ to the subgenome I that is present, respectively, in both polyploids (fig. 2B; supplementary tables S2B, S3). The highly supported subgenomic tree (fig. 2C) recovered the expected phylogeny for the studied diploid *Triticum* and *Aegilops* taxa and demonstrated sister relationships of the homeologous C, F and I subgenomes to their respective *A. speltoides*-like, *A. tauschii-* like and *T. urartu*-like diploid progenitor genomes, which corroborated the accuracy of our subgenome assignment method. According to Marcussen et al. (2014b), our ‘C’ subgenome is equivalent to the subgenome B, ‘F’ to the subgenome D and ‘I’ to the subgenome A, respectively, in the nomenclatural system of the Triticeae genomes.

**Table 1.**
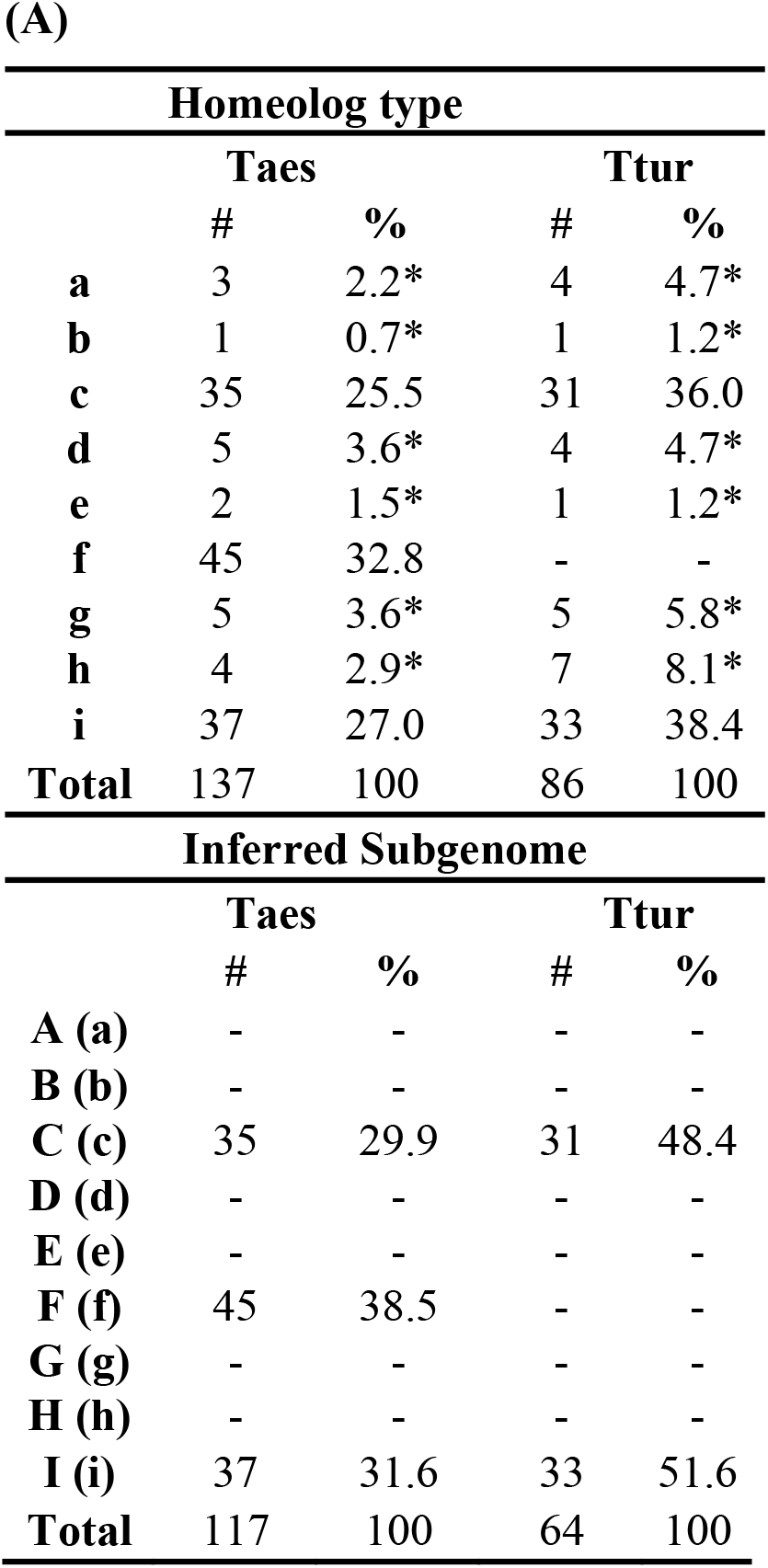
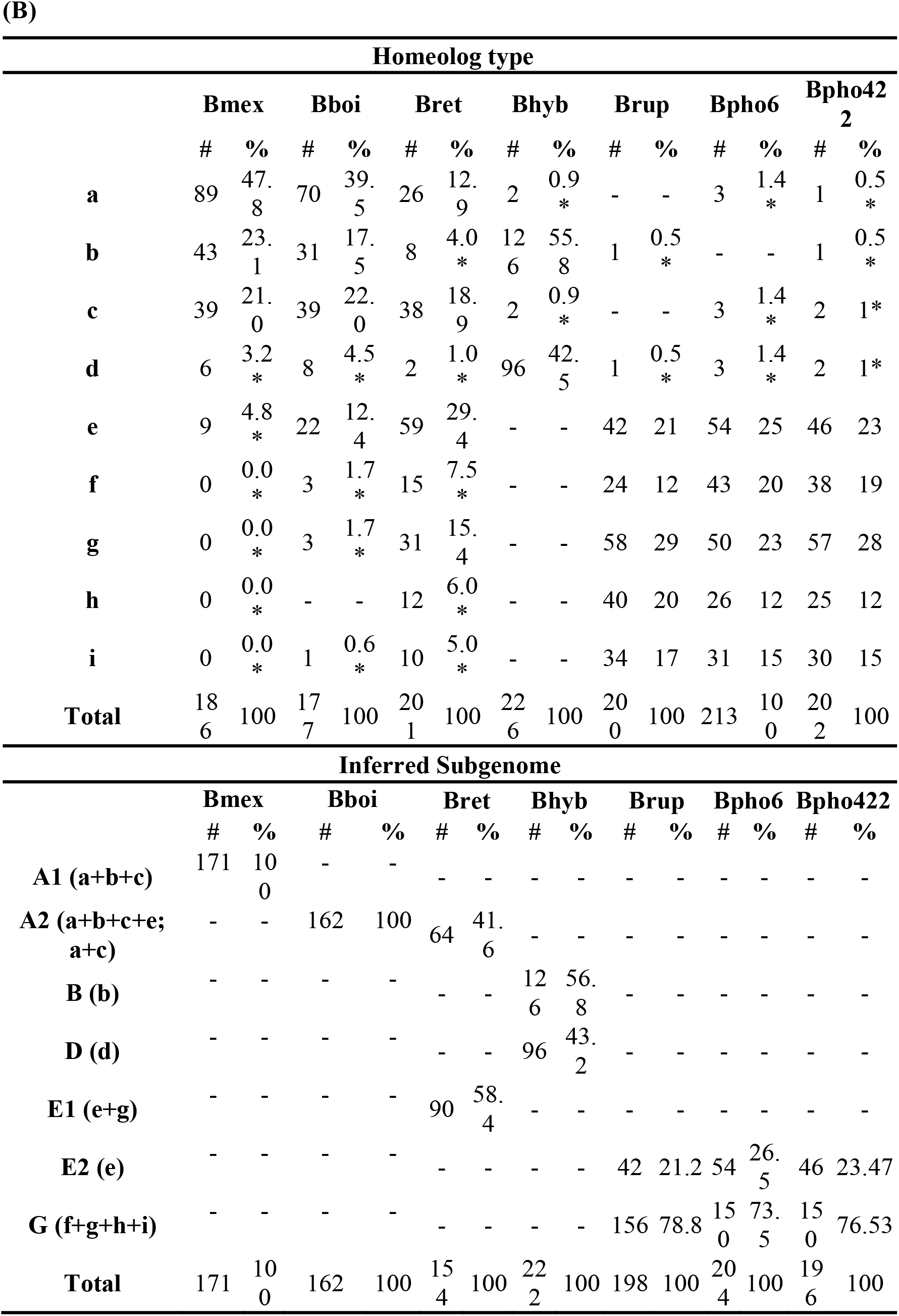
Homeolog allelic and subgenomic data sets. **(A)** *Triticum-Aegilops*. **(B)** *Brachypodium*. Number (#) and percentage (%) of polyploid homeolog alleles that were detected in the studied species by our *Nearest Diploid Species Node* and *Bootstrapping Refinement* algorithms using the aligned genes (a) and core transcripts (b). The homeologs were classified into nine homeolog types (‘a’ to ‘i’) according to their grafting positions in the diploid skeleton tree (figs. 2A, 3A). Those occurring in less than 10% of the selected genes in each accession (see asterisks) were removed from the downstream analyses. The inferred homeologous subgenomes of the studied polyploids that were selected and labeled according to the S*ubgenome Assignment* algorithm and the ploidy level of each polyploid species that was inferred by cytogenetic data. The abbreviations of accessions and cytogenetic data of *Brachypodium* and *Triticum* – *Aegilops* species correspond to those indicated in supplementary tables S1 and S3, respectively.

### Retrieving the known and ‘ghost’ subgenomes of the *Brachypodium* polyploids

The initial *Brachypodium* data set contained 3,675 transcript clusters (supplementary tables S4, S5), which were obtained from a wide transcriptomic analysis of the studied plants (see Methods). These were reduced to 329 MSAs with 1,965 homeologs after the successive filtering steps of the first algorithm (Supplementary Material; supplementary figs. S4A-C). The homeologs were labeled according to the *ad hoc* rules presented in fig. 3A. The bootstrapping refinement algorithm left 322 MSAs and 1,307 homeologs, which were used to build the ML consensus tree (fig. 3B). In contrast to the wheats, *Brachypodium* had an excess of homeolog types that required ranking, selecting the most frequent types and merging some of them in order to retrieve the plausible subgenomes of some polyploids (table 1B; supplementary table S6A). The allotetraploid *B. hybridum* was the exception; it fitted a simple scenario where its ‘b’ and ‘d’ homeologs corresponded to its respective single progenitor subgenomes B (*B. stacei*-type) and D (*B. distachyon*-type). For the remaining *Brachypodium* polyploids, the relatively close ‘a’ and ‘c’ homeolog types (plus ‘b’ in some species) were considered to be variants of the ancestral A subgenomes, the separate ‘e’ homeolog types were assigned to the intermediately evolved E subgenomes and the recently diverged and close ‘f’, ‘g’, ‘h’ and ‘i’ homeolog types were assumed to represent variants of a single core perennial clade G subgenome (figs. 3B, C; supplementary tables S6B, S7). In addition, the clear divergence of the *B. mexicanum* ‘a’ homeolog type from that of *B. boissieri* and *B. retusum* in the ML tree and the PCoA-MST plot (fig. 3B; supplementary fig. S5) supported their respective assignment to the independent ancestral A1 and A2 subgenomes. In contrast, the sequential divergences of the *B. retusum* ‘e’ homeolog type from that of *B. rupestre* and *B. phoenicoides* in the ML tree (fig. 3B) maintained their respective assignments to the independent intermediate E1 and E2 subgenomes (fig. 3C; table 1B; supplementary fig. S5). In this sense and considering the estimated ploidy levels of the studied polyploids (supplementary table S1), the subgenomic assignments were constrained as follows: the tetraploid *B. mexicanum* ‘a’, ‘b’ and ‘c’ homeolog types were assigned to the ancestral A1 subgenome; the hexaploid *B. boissieri* ‘a’, ‘b’, ‘c’ and residual ‘e’ homeolog types to the ancestral A2 subgenome; the tetraploid *B. retusum* ‘a’ and ‘c’ homeolog types to the ancestral A2 subgenome and intermediate ‘e’ (plus recent ‘g’) homeolog types to the intermediate E1 subgenome; the tetraploids *B. rupestre* and *B. phoenicoides* intermediate ‘e’ homeolog types to the intermediate E2 subgenome and the recent core perennial clade ‘f’, ‘g’, ‘h’ and ‘i’ homeolog types to the recent G subgenome (table 1B; supplementary fig. S5; supplementary tables S1, S6, S7).

A strongly supported subgenomic ML consensus tree (fig. 3C), computed from the 322 validated core clusters with single and compound subgenome homeolog types, yielded the same *Brachypodium* gene topology as the dated Bayesian maximum clade credibility (MCC) BEAST tree (fig. 4). The *Brachypodium* stem and crown nodes were estimated to have had Late-Eocene (36.3 Ma) and Mid-Miocene (12.1 Ma) ages, respectively (fig. 4), which is consistent with the previous estimates that were based on a plastome analysis (Sancho et al. 2018). The basic chromosome numbers of the ‘ghost’ and merged subgenomes were inferred from their respective phylogenetic positions and the ploidy levels of the species that contained them (e.g., tetraploid *B. mexicanum* 2n=40: A1 (x=10); hexaploid *B. boissieri* 2n=48: A2 (x=8); allotetraploid *B. retusum* 2n=32: A2 (x=8) and E1 (x=8); allotetraploids *B. rupestre* and *B. phoenicoides* 2n=28: E2 (x=5) and G (x=9) (Figs. 3C, 4; supplementary table S1). The ML tree shows the early divergence of the sister ancestral *B. mexicanum*_A1 (x=10) and *B. boissieri_*A2/*B. retusum*_A2 (x=8) subgenomic ‘ghost’ lineages, which was followed by the successive splits of *B. stacei* (x=10) and its sister derived *B. hybridum_B* subgenomic lineage (x=10) and of *B. distachyon* (x=5) and its sister derived *B. hybridum*_D subgenomic lineage (x=5) (fig. 3C). The split of the ancestral (A1/A2) clade was inferred to have occurred in the Mid-Late-Miocene (10.4 Ma), a time close to that of the split of the oldest extant diploid *B. stacei-type* (genome B) clade (10.2 Ma) (fig. 4). The tree also revealed the successive divergences of the intermediate *B. retusum*_E1 (x=8) and *B. rupestre*_E2/*B. phoenicoides*_E2 (x=5) subgenomic ‘ghost’ lineages and the recently evolved (*B. arbuscula*, (*B. sylvaticum*/*B. pinnatum*)) core perennial clade species (x=9) where the derived *B. rupestre*_G/*B. phoenicoides*_G subgenomic lineages (x=9) were nested within (fig. 3C). The inferred dates indicate that the *B. retusum*_E1 subgenomic ‘ghost’ lineage is more ancestral (4.4 Ma, Early-Pliocene) than the *B. rupestre* and *B. phoenicoides*_E2 subgenome ‘ghost’ lineages (3.8 Ma) (fig. 4). Additionally, the estimated ages for the splits of the core perennial clade (3.0 Ma, Late-Pliocene), the diploid *B. pinnatum/B. sylvaticum* clade (2.1 Ma, Pleistocene) and the *B. rupestre/B. phoenicoides* G subgenomic lineages (2.1 Ma) and the origins of the *B. stacei-type* (2.4 Ma) and *B. distachyon-type* (1.7 Ma) homeologous lineages of *B. hybridum* (fig. 4) are also in agreement with previous datings (Díaz-Pérez et al. 2018; Sancho et al. 2018; Gordon et al. 2020).

**Fig. 4.**
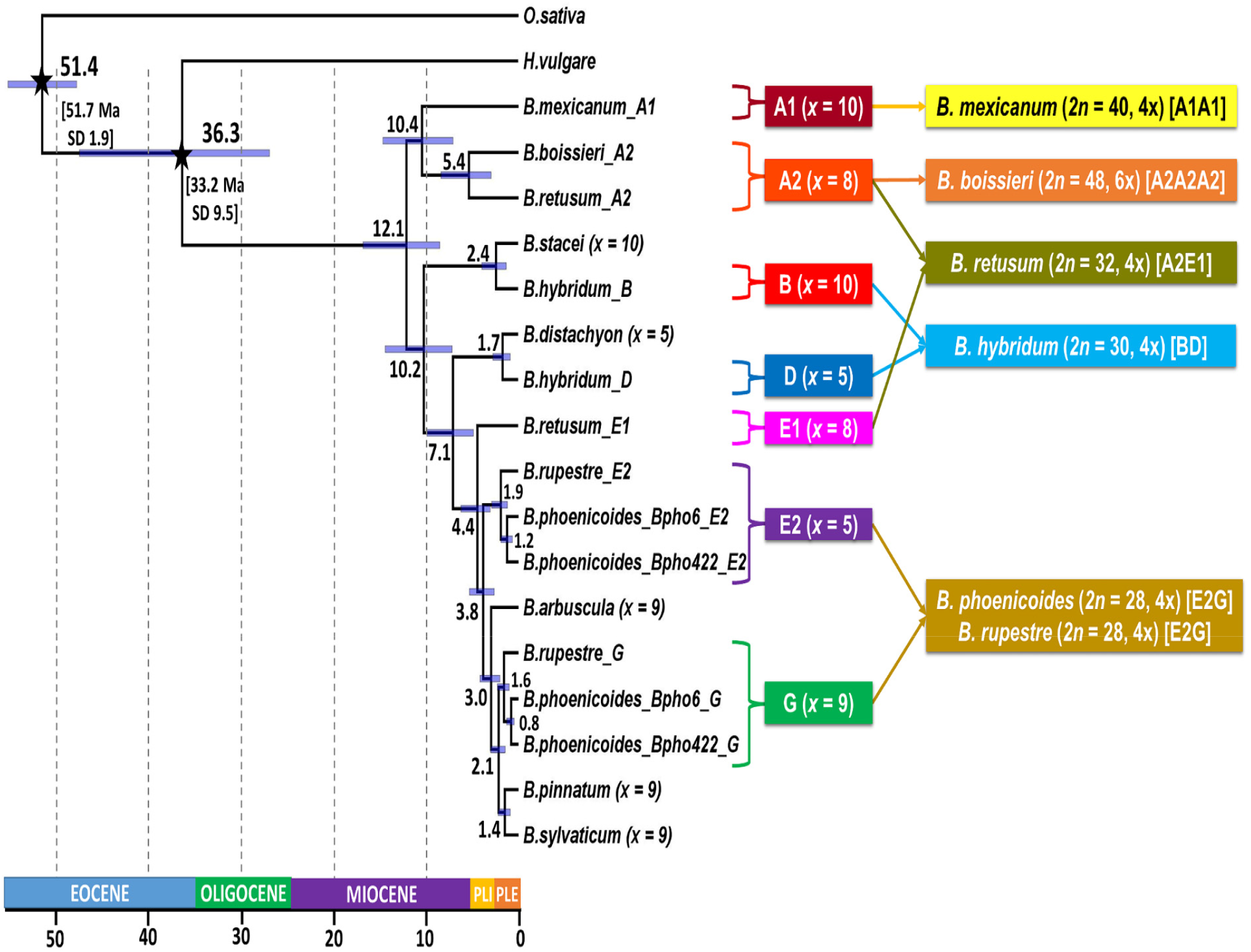
*Brachypodium* Bayesian maximum clade credibility dated chronogram of 322 independent core genes (with the polyploid homeologous subgenomes labeled according to subgenome-types ‘A1’, ‘A2’, ‘B’, ‘D’, ‘E1’, ‘E2’, ‘G’; table 1B) showing the estimated nodal divergence times (medians, in Ma) and the 95% highest posterior density (HPD) intervals (bars). Stars indicate secondary nodal calibration priors (means ± SD, in Mya) for the crown nodes of the BOP [*Oryza* + *Brachypodium* + *Hordeum*] and *Brachypodium* + core pooids [*Brachypodium* + *Hordeum*] clades. Accessions codes of *B. phoenicoides* correspond to those indicated in supplementary table S1.

### Validation of the Subgenome Assignment algorithm using the coalescence methods

The accuracy of our results was validated using a coalescence analysis of the confirmed *Brachypodium* allopolyploid species. We used two strategies: 1) simulated coalescent trees and 2) simulated allopolyploids (fig. 1C; supplementary fig. S1). With Strategy 1 (supplementary tables S8, S9, S10), we evaluated whether the *Subgenome Assignment* algorithm grafted known and ‘ghost’ homeologous subgenomes to the correct branches of the species tree under the hypothetical existence of ILS. Different hypothetical species trees (with variable effective population sizes, see Methods) that contained all of the diploid genomes and one polyploid homeologous subgenome at a time were evaluated with the COAL program (Degnan and Salter 2005), which assays all possible trees that can be constructed with a specific number of tips. In all cases, the highest probability model corresponded to the species tree branch in which the homeologous subgenome was grafted by our *Subgenome Assignment* algorithm (supplementary table S9). There was a complete qualitative agreement between the most frequently observed versus the theoretical topologies for *B. hybridum* [B (b)+D (d)] and for *B. rupestre* and *B. phoenicoides* (Bpho422, Bpho6) [E2+G (g)], which suggests that the theoretical distributions fit closely to the observed data. Similarly, the *B. retusum* [A2 (a)+E1 (e)] and *B. rupestre* [E2 (e)+G (g)] theoretical distribution scenarios had higher frequencies of gene trees in which the A2 subgenome grafted to branch ‘a’ than to ‘c’ and the G subgenome to branch ‘g’ than to ‘h’, respectively, as was scored for the observed data (figs. 3C, 4; supplementary table S9).

Strategy 2 enabled us to measure the ability of the *Subgenome Assignment* algorithm to select the correct allopolyploid subgenomes under different levels of ILS (supplementary tables S11, S12, S13). We generated hypothetical subgenomic allopolyploids that matched the real *Brachypodium* allopolyploids [ancestral-ancestral (A+B), similar to *B. mexicanum* (see Discussion); ancestral-intermediate (A+E), similar to *B. retusum*; and intermediate-recent (E+G), similar to *B. rupestre* and *B. phoenicoides*] and the relative frequency of the theoretical gene tree distributions were calculated for them using COAL. In all cases, the *Subgenome Assignment* algorithm recovered the expected placements of the subgenomes despite the different topological graftings (ancient, intermediate or recent branches) in the diploid skeleton tree at different coalescent-unit levels (0.5 CU, 1 CU) of ILS (supplementary table S12). This suggests that our algorithm is able to place the subgenomes in their correct branch independently of any deep or shallow coalescences (branch lengths) or the effective population sizes of the *Brachypodium* lineages.

### Karyotypic identification of the new *Brachypodium* genomes using comparative chromosome barcoding

The karyotypes of the previously unstudied *B. arbuscula* (2n=18), *B. boissieri* (2n=48) and *B. retusum* (2n=32) species were analyzed using CCB mapping as was described in Lusinska et al. (2019). The mapping was done with reference to the *B. distachyon* karyotype and its genome was compared to the ancestral rice genome (IBI, 2010). The arrangement of all of the BAC clones is shown on the cytogenetic maps of *B. arbuscula* (fig. 5A), *B. boissieri* (fig. 5B) and *B. retusum* (fig. 5C) chromosomes.

**Fig. 5.**
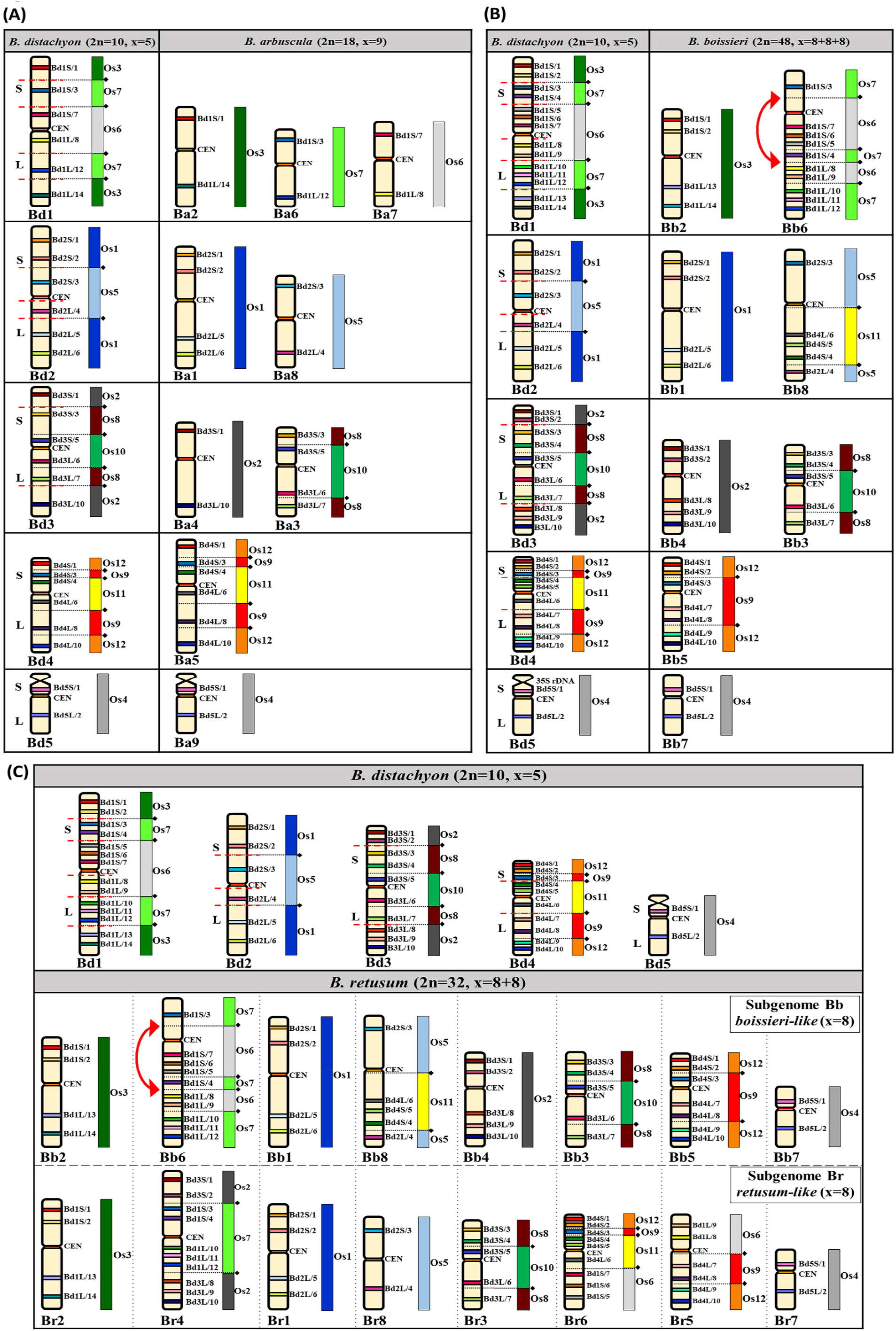
Distribution of the BAC clones derived from chromosomes Bd1-Bd5 of *B. distachyon* (2n=10, x=5) that were comparatively mapped to, respectively, the chromosomes of the (A) diploid *B. arbuscula* (2n=18, x=9), (B) autohexaploid *B. boissieri* (2n=48, x=8+8+8) and (C) allotetraploid *B. retusum* (2n=32, x=8+8). Only one homologue from a pair is shown. The diagrams next to the *Brachypodium* (Bd, Ba (A); Bd, Bb (B); Bd, Bb, Br (C)) chromosomes align the BAC clones to the homeologous regions (syntenic segments) in the relevant ancestral rice chromosome equivalents Os1-Os12. Black diamonds and dotted lines indicate the hypothetical fusion points of the ancestral rice chromosome equivalents (adapted from IBI 2010). Red dashed lines indicate the chromosomal breakpoints in the Ba-genome chromosomes of *B. arbuscula* (A), Bb-genome chromosomes of *B. boissieri* (B) and Bb- and Br-subgenome chromosomes in *B. retusum* (C) that were found using CCB. Red arrows point to a pericentric inversion that was found on chromosome Bb6 (B, C).

The cytomolecular mapping of the diploid *B. arbuscula* showed that each of the BAC clones hybridized to a single chromosome pair (fig. 5A; supplementary figs. S6-S7). The karyotypic pattern of this species was revealed to be the same as that of the genomes of the core perennial clade diploids *B. sylvaticum* and *B. pinnatum* with x=9 (fig. 6; Lusinska et al. 2019). *B. arbuscula* chromosomes Ba1, Ba2, Ba4, Ba6, Ba7, Ba8 and Ba9, which correspond, respectively, to the ancestral *Oryza sativa* Os1, Os3, Os2, Os7, Os6, Os5 and Os4 chromosomes did not undergo any fusions, whereas one nested chromosome fusion (NCF) of Os8 and Os10 was observed on chromosome Ba3 and two NCFs of Os12+Os9+Os11 on chromosome Ba5 (fig. 5A; supplementary figs. S6-S7).

**Fig. 6.**
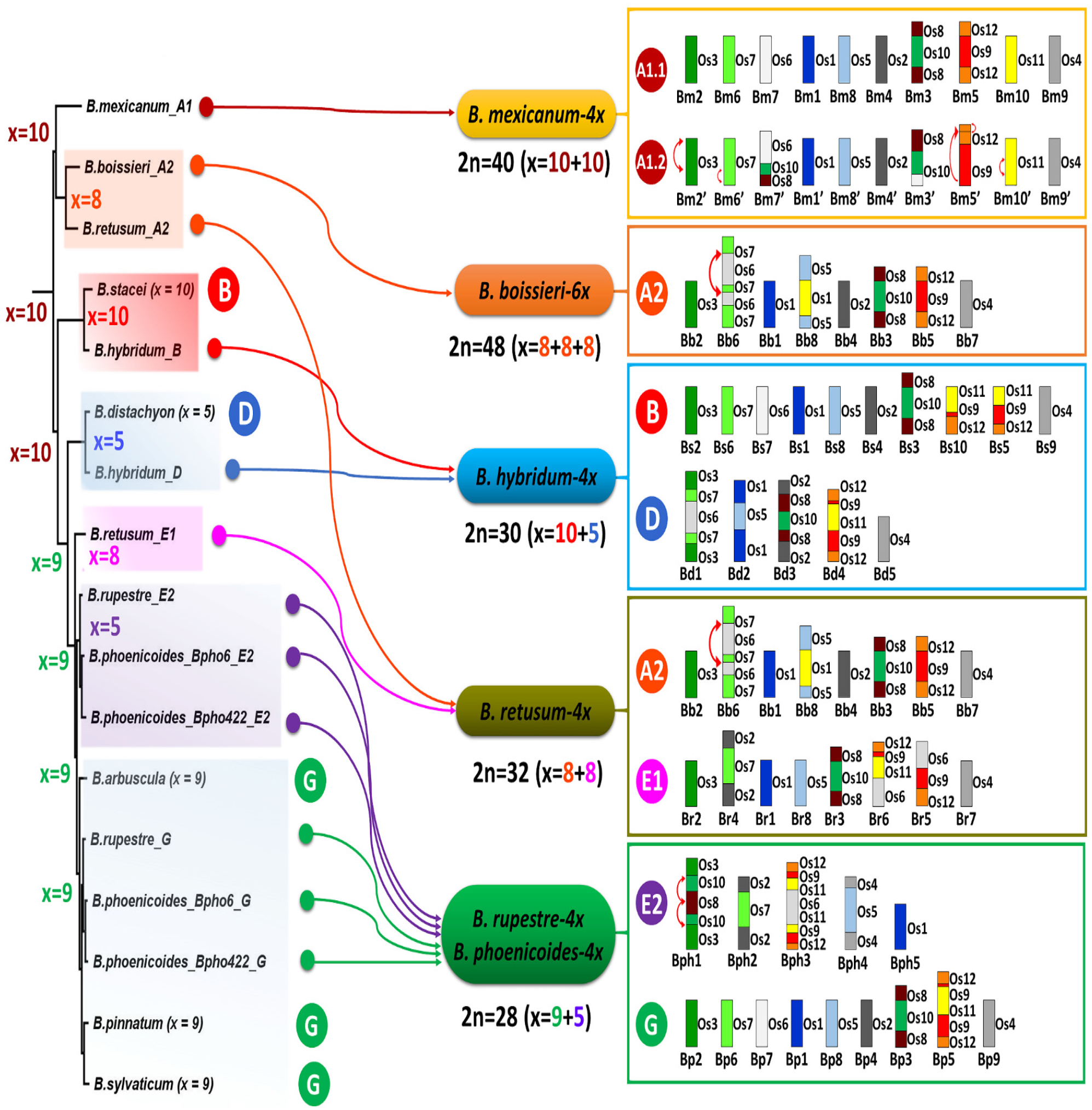
A comprehensive evolutionary framework for the origin of *Brachypodium* allopolyploids based on the combined phylogenomic and comparative chromosome barcoding analyses. Colors indicate the different types of (sub)genomes that were retrieved in the phylogenomic analysis and letters designate the karyotype profiles that were found in the diploids and polyploids. The arrows link the inferred (sub)genomes and karyotypes of each studied *Brachypodium* polyploid. The karyotype models are based on the CCB analysis of the *B. arbuscula* (2x), *B. retusum* (4x) and *B. boissieri* (6x) species that were analyzed in this study and other *Brachypodium* representatives that had been previously studied (Lusinska et al. 2018; Lusinska et al. 2019; Gordon et al. 2020). Within the karyotypes, each chromosome or homoeologous chromosome region corresponded to the relevant ancestral rice chromosome equivalents (Os1-Os12; Os – *Oryza sativa*) (IBI 2010). The basic chromosome numbers (x) that were obtained for each genome and karyotype and inferred for the ancestors of the subgenomic tree are shown in the topology; their colors correspond to their respective (sub)genomic and karyotypic assignments. (Sub)genome designations: ‘A1’ – ancestral *B. mexicanum* (dark red), ‘A2’ – ancestral *B. boissiei* (orange), ‘B’ – *B. stacei* (red), ‘D’ – *B. distachyon* (blue), ‘E1’ – intermediate *B. retusum* (purple), ‘E2’ – intermediate *Brachypodium* core perennials 4x (violet), ‘G’ – recent *Brachypodium* core perennials 4x (green) (table 1). Chromosome designations within the (sub)genomes: Bb – *B. boissiei*, Bd – *B. distachyon*, Bm, Bm’ – *B. mexicanum*, Bp – *Brachypodium* core perennials x=9, Bph – *Brachypodium* core perennials x=5, Br – *B. retusum*, Bs – *B. stacei*.

The CCB of the *B. boissieri* (2n=48) chromosomes revealed that each BAC had six hybridization sites that were localized on three chromosome pairs (fig. 5B; supplementary figs. S8-S12). The identical triplicated distribution pattern of the BAC-FISH signals in the morphologically uniform chromosomes supports the autohexaploid nature of *B. boissieri* with x=8 (figs. 5B, 6). The CCB analysis also detected the presence of chromosome fusions and rearrangements that were specific for the *B. boissieri* genome. *B. boissieri* chromosomes Bb1, Bb2, Bb4 and Bb7 correspond, respectively, to the Os1, Os3, Os2 and Os4 chromosomes, whereas Bb3 resulted from the NCF of Os8 and Os10, Bb5 from the NCF of Os12 and Os9, Bb6 from the NCF of Os7 and Os6 that were complemented with pericentromeric inversion and Bb8 from the NCF of Os5 and Os11, which is a unique trait of the *B. boissieri* karyotype (figs. 5B, 6; supplementary figs. S8-S12). In addition, the *B. boissieri* genome is the only one in *Brachypodium*, other than that of *B. mexicanum*, that does not have Os12+Os9 fused with Os11 (fig. 6; Lusinska et al. 2019).

The BAC-FISH analysis of *B. retusum* (2n=32) demonstrated that each clone hybridized to four sites that were located on two chromosome pairs (fig. 5C; supplementary figs. S13-S16). Unlike *B. boissieri*, this species had two distinct groups of chromosomes, each consisting of eight pairs of chromosomes, thereby revealing an allotetraploid nature for this *B. retusum* cytotype (figs. 5C, 6). One of the chromosomal sets corresponded to a subgenome with the same karyotypic pattern as that of the *B. boissieri* genome and the other to a new subgenome with a *B. retusum-type* karyotype, both with x=8 (figs. 5C, 6; supplementary figs. S13-S16). The *B. retusum* chromosomes Br1, Br2, Br7 and Br8 corresponded, respectively, to Os1, Os3, Os4 and Os5 chromosomes, whereas Br3 resulted from the NCF of Os8 and Os10 and Br4 from the NCF of Os2 and Os7 (figs. 5C, 6), which is a trait that was shared with the ‘ghost’ subgenome x=5 present in the core perennial clade allotetraploids with 2n=28 (fig. 6; Lusinska et al. 2019). The *B. distachyon* Bd1- and Bd4-BAC-derived probes hybridized to two different *B. retusum* Br5 and Br6 chromosomes, both of which are specific to this subgenome in their syntenic segment composition. The distinctive arrangement of the BAC-FISH signals indicates that these two chromosomes originated *via* the reciprocal translocation of two ancestral *Brachypodium* chromosomes that correspond to Os12+Os9+Os11 and Os6 (figs. 5C, 6; supplementary figs. S13-S16; Lusinska et al. 2019).

## Discussion

Deciphering the diploid origins of allopolyploids faces the challenge of accurately capturing their progenitor subgenomes (Levin 2013; Bombarely et al. 2014; Soltis et al. 2016). Approaches using the coalescent-based analyses of multi-labeled trees and networks are hindered by homeolog loss and ILS (Marcussen et al. 2015; Thomas et al. 2017). The deconvolution of hybrid subgenomes is still challenging, especially in the absence of any known extant parents and of whole genome sequence data for the studied species (Soltis et al. 2016; Liston et al. 2020). Recently, a phylogenetic subgenome-tree searching (PhyDS) pipeline was developed to retrieve the four progenitor diploid genomes of allo-octoploid *Fragaria x ananassa* from a wide transcriptome analysis of candidate species (Edger et al. 2019). This method explored the exclusive clades that contained the syntenic ortholog and homeolog sequences and identified the progenitor subgenomes based on bootstrap support cut-off values across the genes (Edger et al. 2019; Edger et al. 2020). There is current debate, however, on the accuracy of this approach, which although it correctly identified two extant progenitor diploid species (*F. vesca*, *F. iinumae*), it may have failed to identify the other two (Liston et al. 2020; Feng et al. 2021; Session and Rokhsar 2020). Feng et al. (2021), when mapping the *Fragaria x ananassa* genomic reads to five diploid *Fragaria* genomes and comparing the ortholog between the cultivated strawberry and the five diploid species, recognized *F. vesca* and *F. iinumae* as being progenitors of *Fragaria x ananassa*, but rejected the participation of *F. viridis* and the other studied diploids in the origin of the octoploid. Similar findings were obtained by Session and Rokhsar (2020) *via* a chromosomal distribution analysis of the transposable elements in the *Fragaria x ananassa* genome. However, both studies were unable to identify the other two ‘ghost’ subgenomes because of their omission of an algorithm to take into account the potential existence of unknown progenitors.

Our PhyloSD pipeline refines the previous methods and additionally enables the progenitor genomes of an unknown origin in the polyploids to be inferred. This was accomplished by filtering the gene trees that were congruent with the strongly supported diploid skeleton tree, the use of a close outgroup that would enable the detection of ‘ghost’ genomes that had diverged prior to the divergence of extant diploid genomes, the grafting of most common homeolog sequences to the nodal-branch groups of the diploid skeleton tree and their assignment to the ploidy-level and CCB-informed main subgenomes. Our method was validated in the thoroughly studied *Triticum*-*Aegilops* polyploid complex in which the homeologous subgenomes of the *T. turgidum* 4x and *T. aestivum* 6x allopolyploids (Marcussen et al. 2014a) were accurately inferred and in the allotetraploid *Brachypodium hybridum*, which was used as the internal control species with known progenitor genomes (Catalán et al. 2012; Gordon et al. 2020). In *Brachypodium*, our strategy enabled us to uncover three known (B, D and amalgamated G) and four unknown (A1, A2, E1, E2) diploid progenitor genomes of six polyploid *Brachypodium* species that had different dysploid ancestral origins (fig. 3; supplementary fig. S5). Moreover, the inferences of the *Subgenome Assignment* algorithm were robust to the presence of the ‘ghost’ subgenomes and to the moderate existence of ILS in *Brachypodium* (Strategy 1, supplementary table S9; Strategy 2, supplementary table S12).

One of the caveats of our approach is that only a small percentage of the pre-filtered gene clusters had a topology that was congruent with that of the diploid species tree (18% in *Triticum-Aegilops;* 17% in *Brachypodium*) and only those genes could be used to infer the homeologous subgenomes of the allopolyploids. Although other phylogenetic approaches such as PhyDS also use low percentages of the total number of expressed genes (<17.9%) to identify the progenitor subgenomes of allopolyploids, in this case, this was done by selecting the high-confidence syntenic homeologs that are present in each of the subgenomes of the polyploid (Edger et al. 2019; Edger et al. 2020). Recent approaches have proposed the inclusion of paralogs in order to increase the amount of data that can be used to infer a species tree (Smith and Hahn 2020). However, the all-gene total evidence principle could lead to misleading phylogenies if the increasing amount of data also increases the phylogenetic noise. By contrast, the utility of our PhyloSD pipeline concurs with a restricted total evidence genomic scenario that favors the use of selected components of the data partitions that better fit the evolutionary models as the most reliable method for phylogenetic reconstruction (Goremykin et al. 2015) and, consequently, for homeolog subgenomic detection.

The hypothetical genomes A1 of *B. mexicanum*, A2 of *B. boissieri* and *B. retusum*, E1 of *B. retusum* and E2 and G of *B. phoenicoides* and *B. rupestre* are partially similar to those that were retrieved using a few cloned nuclear ribosomal genes (Catalán et al. 2016; Díaz-Pérez et al. 2018); however, here, they are supported by a larger set of 322 core-expressed genes (fig. 3C; table 1B; supplementary table S14). Our phylogenomic results were independently confirmed by our comparative chromosome barcoding data (Figs. 5, 6; Supplementary Figs. S6-S16; Lusinska et al. 2019). Our CCB karyotypes undisputedly identified the three known, and four ‘ghost’ diploid progenitor genomes that are present in the six studied *Brachypodium* polyploids (fig. 6). The feasibility of our approach was facilitated by the high synteny that was observed across the *Brachypodium* reference genomes (Scholthof et al. 2018; Gordon et al. 2020) and by the high integrity of the progenitor genomes that were found in the subgenomes of some of the *Brachypodium* allopolyploids (Gordon et al. 2020). The *Brachypodium* genomes likely derived from a karyotype evolution model of successive centromeric chromosome fusions with a relatively low incidence of other types of rearrangements (fig. 6; Lusinska et al. 2019).

Although our subgenome detection algorithms were initially designed to retrieve the homeologous subgenomes of known and putative allopolyploids, two of the studied *Brachypodium* polyploids revealed homeolog types that could be assigned to the compound (sub)genomes that pertain to autopolyploids (*B. mexicanum* A1A1, *B. boissieri* A2A2A2) (figs. 3, 4). These findings were confirmed by our CCB data (figs. 5, 6; supplementary figs. S8-S12; Lusinska et al. 2019). By contrast, all of the remaining *Brachypodium* polyploid species were undisputedly identified as allopolyploids (figs. 3C, 4, 5, 6; supplementary figs. S13-S16; table 1B; Lusinska et al. 2019). The relatively large genome size of *B. mexicanum*, which is not found in other *Brachypodium* species (supplementary table S1) and the uncertainty in the assignment of its close ancestral homeologs to a single genome (A1) or to two closely related genomes [e.g., A1.1 (‘a’+’c’) and A1.2 (‘b’)], would also favor an alternative segmental allopolyploid scenario (Mason and Wendel 2020) for this species. In fact, this agrees with the similar karyotypic barcoding patterns that were observed in its two chromosome complements (fig. 6; Lusinska et al. 2019). The minor intrachromosomal inversions and translocations that were detected in Bm2, Bm5, Bm6 and Bm10 and the reciprocal translocation of Bm7 and Bm3 between the two chromosomal sets (fig. 6; Lusinska et al. 2019) could have been inherited from close but independent diploid progenitors or could have resulted from post-polyploidization rearrangements of a duplicated autotetraploid. This would support the alternative hypothetical scenarios of segmental allotetraploidy vs. autotetraploidy, which would require more precise genomic and cytomolecular data to be tested. *B. boissieri* was revealed to be an unequivocal autohexaploid (figs. 3, 5B, 6). The assignment of its close ‘a’, ‘b’ and ‘c’ plus the residual ‘e’ homeologs to a unique A2 x=8 genome (Table 1B; fig. 3) was corroborated by its robust and irrefutable karyotype (figs. 5B, 6; supplementary figs. S8-S12). To date, this species constitutes the only supported evidence of autopolyploidy within *Brachypodium*.

The successive divergences of the *Brachypodium* polyploid subgenomes and their karyotype structures support an evolutionarily descendant dysploidy trend from the ancestral x=10 (A1, B) to the recent x=9 (G) genomes (figs. 4, 6), which corroborates the findings of Lusinska et al. (2019) that inferred the existence of an ancestral *Brachypodium* karyotype (ABK) of x=10. Our newly emerged karyotype evolutionary scenario of *Brachypodium* also involves two independent reductions from x=10 to x=8 (ancestral A2) and from x=9 to x=8 (intermediate E1) plus two independent reductions from x=9 to x=5 (intermediate D and E2) (figs. 4, 6). The nearly contemporary Mid-Late Miocene inferred origins of the divergent A1 and B x=10 genomes (fig. 4) resulted in highly syntenic karyotypes that only had rearrangements within some of the homeologous chromosomes (e.g., chromosomes Bm5 and Bm10 of subgenome A1.1 and intrachromosomal rearrangements in Bm5’ and Bm10’ of subgenome A1.2 of *B. mexicanum* vs. chromosomes Bs5 and Bs10 of *B. stacei* that probably originated *via* a reciprocal translocation or chromosome split) (fig. 6). By contrast, the parallel but separate reductions to x=8 and x=5 genomes implies major structural changes that primarily affected the number and compositions of the chromosomal fusions. Thus, the two hypothesized NCFs that resulted in the ancestral Late-Miocene A2 x=8 genome (chromosomes Bb6 and Bb8) differed from the more complex pattern of the three NCFs plus one translocation that resulted in the Early-Pliocene E1 x=8 genome (chromosomes Br4, Br5 and Br6) (figs. 4, 6). Similarly, the increasing reduction that was caused by the four NCFs from the hypothetical x=9 Intermediate Ancestral Brachypodium Karyotype (ABK) (fig. 6; Lusinska et al. 2019) that ended in the Late-Miocene D x=5 genome (chromosomes Bd1 to Bd4) was distinct from the four NCFs that resulted in the Late-Pliocene E2 x=5 genome (chromosomes Bph1 to Bph4) (figs. 4, 6). Despite the large hypothesized rearrangements that were experienced by the *Brachypodium* genomes (fig. 6), their chromosomes are highly collinear as is demonstrated by the high synteny that was observed between the reference genomes of the ancestral *B. stacei* x=10 (B) genome and the intermediate and highly reduced *B. distachyon* x=5 (D) genome (Gordon et al. 2020). These findings, together with the inferred ages and karyotype patterns (figs. 4, 6), support a highly dynamic evolutionary scenario of chromosomal reshufflings that led to diploid species that have highly syntenic but rearranged genomes during the last 12 million years. Interestingly, the diploid progenitor genomes have remained almost intact in the derived allopolyploid subgenomes as is demonstrated in the inherited karyotypes and collinear sequences of the allotetraploid *B. hybridum* reference subgenomes and those of its diploid progenitors’ reference genomes (fig.. 6; Gordon et al. 2020). These genomic evidence supports our assumption that the identified ‘ghost’ genomes of the *Brachypodium* polyploids (A1, A2, E1, E2; figs. 3, 4, 6) are the preserved vestiges of the diploid progenitor genomes they had originated from.

While our current analyses suggest that the ancestral A1 and A2 and the intermediate E1 and E2 genomes likely correspond to extinct or unsampled diploid *Brachypodium* species, identifying the G genome is more problematic. The phylogenomic data could not accurately assign the very close ‘f’, ‘g’, ‘h’ and ‘i’ homeologs to any of the core perennial clade diploid lineages (table 1B; fig. 3) and the CCB data could not detect differences in the karyotypic patterns of *B. arbuscula* (figs. 5A, 6), *B. sylvaticum* and *B. pinnatum* and the x=9 subgenomes of the allotetraploids *B. phoenicoides* and *B. rupestre* (fig. 6; Lusinska et al. 2019). The characterization of the Plio-Pleistocene-originating core perennial *Brachypodium* diploids would require the use of a large number of highly variable genomic loci and chromosomal barcodes. Still, the accurate identification of the four ‘ghost’ genomes by our combined PhyloSD and CCB methods makes *Brachypodium* a unique case within the angiosperms. This model genus constitutes an excellent study system to investigate the impact of the ‘ghost’ subgenomes on the functional, adaptive and evolutionary behavior of their hosting polyploids.

## Conclusions

Our results demonstrate the value of the PhyloSD pipeline coupled with the CCB approach in detecting polyploid subgenomes. The wheat benchmark indicated that it can identify the diploid homeologous subgenomes from extant progenitors. More importantly, it also identified three known and four novel ‘ghost’ subgenomes in *Brachypodium*, thus shedding light on the complex and intricate evolutionary history of this grass model genus. Our method could be of significant value in studies of polyploid plants that have complex histories of hybridizations and polyploidizations.

## Material and Methods

### Sampling, chromosome counting and genome size determination

Eleven *Brachypodium* species and two ecotypes [all main diploids *B. arbuscula, B. distachyon*, *B. pinnatum*, *B. stacei* and *B. sylvaticum* and polyploids *B. boissieri, B. hybridum, B. mexicanum, B. phoenicoides* (Bpho6 and Bpho422 accessions), *B. retusum* and *B. rupestre*] were studied (supplementary table S1). The genome size (GS) and chromosome counting estimations were performed using flow cytometry and on DAPI-stained meristematic root cells following the protocols of Doležel et al. (2007) and Jenkins and Hasterok (2007), respectively. The ploidy levels were inferred from the chromosome counts (2n) and the GS (pg/2C) estimations that were performed in the same accessions that were used in the transcriptome study and through the GS and 2n values that were obtained in conspecific accessions that showed similar values (supplementary table S1).

### Transcriptomic data of *Brachypodium*

Total RNA was isolated from the leaf tissue of each individual plant under one of the following conditions: control, soil-drying stress, heat stress and salt stress; pooled RNAs were used for the sequencing. The transcript sequences were assembled using trinityrnaseq-r20140717 (Grabherr et al. 2011). A *de novo* assembly of the *Brachypodium* RNA-seq reads (supplementary table S4) produced 72 to 160 thousand transcript isoforms with median lengths ranging between 414 to 555 bp (supplementary table S5). The *Brachypodium* RNA-seq data were deposited in the ENA (European Nucleotide Archive; https://www.ebi.ac.uk/ena; see supplementary Supplemental Methods). The RNA-seq data of *B. distachyon* (Bd21) and *B. sylvaticum* (Brasy-Esp) were obtained from Bettgenhaeuser et al. (2017) and Fox et al. (2013), respectively, and data of the outgroups *Oryza sativa* (SRX738077) and *Hordeum vulgare* (ERR159679) were obtained from the INSDC archives.

### Genomic data of *Triticum-Aegilops*

The genomic sequence data of the six *Triticum* and *Aegilops* species were retrieved from Marcussen et al. (2014b). The cDNA sequences of *T. turgidum* were retrieved from Maccaferri et al. (2019) and those of the outgroups *O. sativa, B. distachyon* and *H. vulgare* were retrieved from Ouyang et al. (2007), IBI (2010) and Mascher et al. (2017), respectively.

### Phylogenomic and dating analyses

The *Brachypodium* and *Triticum*-*Aegilops* data sets were aligned using GET_HOMOLOGUES-EST v09112017 (Contreras-Moreira et al. 2017) and MAFFT v7.222 (Katoh et al. 2002; Katoh and Standley 2013), respectively. The ML analyses were performed using IQ-TREE v.1.6.1 (Minh et al. 2013; Nguyen et al. 2014; Chernomor et al. 2016; Kalyaanamoorthy et al. 2017). The ultrafast bootstrap searches were replicated 1000x. The distance-based coalescence analyses of the *Brachypodium* and *Triticum-Aegilops* diploid MSAs were performed using ASTRAL v5.7.3 (Zhang et al. 2018) and STAR and STEAC (R v.3.5.1; Liu and Yu 2010). The Bayesian phylogenetic dating analysis of the *Brachypodium* data set was conducted using BEAST 2.4.7 (Bouckaert et al. 2014).

### Performance of the Subgenome Assignment algorithm in the presence of ILS in *Brachypodium*

The COAL program (Degnan and Salter 2005) was used to compute the theoretical probabilities of the gene tree topologies from fixed species trees under a multispecies ILS scenario using two strategies (supplementary tables S8-S13). The *Subgenome Assignment* algorithm was applied to each set of probabilities that had been computed from COAL for a single species tree and the selected subgenomes were matched to the expected subgenomes in order to validate the algorithm. In Strategy 1, the homeologous subgenomes of the *Brachypodium* allopolyploids were coded as for the observed data and the divergence time for each polyploid lineage was inferred from the closer ancestral node that included it as a sister lineage to its diploid species (figs. 3C, 4; supplementary tables S8-S10). This species tree was used to compute the theoretical distribution of all of the gene tree topologies that had the optimal diploid skeleton topology and that contained the polyploid subgenome. The branch lengths were transformed to Coalescence Units (CU), where CU=g/2Ne, and assuming g=1.5 years per generation and optimal effective population sizes of Ne=5E5, Ne=1E6 and Ne=2E6. The theoretical distributions from two homeologous subgenomes were proportionally merged in order to recreate the genomic compositions of the *Brachypodium* allotetraploids, following the same criteria as for the observed homeolog types (fig. 3B, C). In Strategy 2, an effective population size of Ne=5E5 individuals and two branch lengths were tested for all of the lineages of a fixed tree using different coalescent units [deep coalescence (1 CU), equivalent to 1.5 My; and shallow coalescence (0.5 CU), equivalent to 0.75 My (supplementary tables S11-S13)].

### Comparative chromosome barcoding

Three unsurveyed perennial *Brachypodium* species, *B. arbuscula* Barb502 (2n=2x=18), *B. boissieri* Bbois10 (2n=6x=48) and *B. retusum* Bret504 (2n=4x=32), were analyzed in this study together with the reference *B. distachyon* Bd21 (supplementary table S1). A multisubstrate chromosome preparations (reference *B. distachyon* plus another *Brachypodium* species at a time) were prepared of the root-tip meristems as was described in Hasterok et al. (2006). The 43 BAC clones (supplementary table S15) that were used in this study had previously been employed in the construction of the karyotypes of the other *Brachypodium* genomes (fig. 6; Lusinska et al. 2019). These probes came from the BD_ABa and BD_CBa genomic DNA libraries that had been generated from the five assemblies of FingerPrinted Contigs that had been assigned to the respective reference chromosomes of *B. distachyon* (Febrer et al. 2010). In order to determine any potential intraspecific variation, each clone was mapped to the chromosome preparations of at least three individuals of each species or accession. The probe labeling with nick translation using tetramethylrhodamine-5-dUTP, digoxigenin-11-dUTP or biotin-16-dUTP (all Sigma-Aldrich) and FISH were performed according to Jenkins and Hasterok (2007) with minor modifications (Lusinska et al. 2018). The images were acquired using a wide-field epifluorescence microscope (AxioImager.Z.2, Zeiss) and a high-sensitivity monochromatic camera (AxioCam Mrm, Zeiss) and then uniformly processed using ZEN 2.3 Pro (Zeiss) and Photoshop CS3 (Adobe).

## Supplementary material

The supplementary information that accompanies this paper can be found at Dryad repository https://doi.org/10.5061/dryad.ncjsxksqw (temporary dryad link: https://datadryad.org/stash/share/TqyetD7bwKsxb1jRW7Sz9wRZwb63VBPvKBSoWTrrcyU).

Additional information on *Brachypodium* sampling, cytogenetic and transcriptomic studies, *Triticum*-*Aegilops* genomic data, phylogenomic and dating analysis and performance of the *Subgenome Assignment* algorithm under coalescence scenarios is indicated in Supplementary Material.

## Availability of data and materials

The complete PhyloSD protocol (source code, step-by-step instructions, commands and examples) is available in Github (https://github.com/eead-csic-compbio/allopolyploids).

The supplementary information (supplementary tables and figures and Supplementary Material), detailed pipelines and algorithms, alignments, BEAST xml file and phylograms of the *Brachypodium* and *Triticum-Aegilops* groups are available in Dryad. (https://doi.org/10.5061/dryad.ncjsxksqw; temporary Dryad link: https://datadryad.org/stash/share/TqyetD7bwKsxb1jRW7Sz9wRZwb63VBPvKBSoWTrrcyU).

## Author contributions

PC, RS, BC, AD and RH designed the study. RS, LI and JL developed the experimental work. RS, BC, AD, LI, JL, RH and PC analyzed the data and interpreted the results. RS, AD, BC and PC prepared the manuscript. All of the authors revised the manuscript.

## Conflict of interest/Competing interests

The authors declare that there are no conflicts of interests/competing interests.

## Funding

This work was supported by the Spanish Ministries of Economy and Competitivity (Mineco) and Science and Innovation (MICINN) CGL2016-79790-P and PID2019-108195GB-I00 and University of Zaragoza UZ2016_TEC02 grant projects and funding from Harvard University. The work conducted by the US DOE Joint Genome Institute was supported by the Office of Science of the US Department of Energy (DOE) under Contract no. DE-AC02-05CH11231. RS was funded by a Mineco FPI PhD fellowship, Mineco and Ibercaja-CAI mobility grants and Instituto de Estudios Altoaragoneses grant. BCM was funded by Fundación ARAID. BCM, PC and RS were also funded by a European Social Fund/Aragón Government Bioflora research grants A01-17R and A01-20R. The work that was conducted at the University of Silesia in Katowice was supported by the Research Excellence Initiative program.

## Acknowledgments

We would like to thank the following institutions for providing the facilities to develop this study. The *Brachypodium* plants that were used in the study were collected in the field and cultivated at the High Polytechnic School of Huesca (University of Zaragoza) and vouchered specimens were deposited in the JACA (Pyrenean Institute of Ecology-CSIC) and UZ (University of Zaragoza) herbaria. The propagated clones were imported to and grown at The Arnold Arboretum of Harvard University under USDA APHIS PPQ PCIP-16-0433. The RNA-seq laboratory analyses were conducted at the Universities of Zaragoza and Harvard, the cytomolecular analyses at the University of Silesia in Katowice and the bioinformatic and phylogenomic analyses at the EEAD-CSIC and University of Zaragoza.

## Notes

https://github.com/eead-csic-compbio/allopolyploids

https://datadryad.org/stash/share/TqyetD7bwKsxb1jRW7Sz9wRZwb63VBPvKBSoWTrrcyU

### Competing Interest Statement

The authors have declared no competing interest.

