## Supplementary_Materials for "Tracking the Ancestry of Known and ‘Ghost’ Homeologous Subgenomes in Model Grass *Brachypodium* Polyploids"

**Supplementary material**

**Development of the PhyloSD pipeline. Step-by-step application of the algorithms to the benchmarked *Triticum*-*Aegilops* data set and the *Brachypodium* case study*.* Each step is numbered and referenced to the bioinformatic workflow shown in Supplementary figure S1.**

**1. ‘Nearest Diploid Species Node’ algorithm**

***1.1. Filtering multiple sequence alignments (MSA) and defining the diploid compact block of sequences***

In *Triticum*-*Aegilops*, the starting alignment consisted of a data set of 275 ortholog clusters of five diploid [*Aegilops sharonensis* (Asha)*, A. speltoides* (Aspe)*, A. tauschii* (Atau)*, Triticum monococcum* (Tmon) and *T. urartu* (Tura)], two polyploid [*T. aestivum* (Taes) 6x and *T. turgidum* (Ttur) 4x] from Marcussen et al. (2014b) and Maccaferri et al. (2019) and three diploid outgroup [*B. distachyon* (Bdis)*, H. vulgare* (Hvul) and *O. sativa* (Osat)] samples from Ouyang et al. (2007), IBI (2010) and Mascher et al. (2017), respectively (see References in main manuscript). The *trim_MSA_block* tool was executed to obtain a compact block of unique sequences for each diploid species, removing short sequences (<100 bp). Alignments that did not include the diploid species were removed. Polyploid sequences that did not overlap at least 50% of the diploid block sequences’ lengths were discarded. The newly obtained MSAs were further processed by trimAl v1.4.rev15 Capella-Gutiérrez et al. (2009) using -automated1 option to remove spurious sequences or poorly aligned regions, resulting in 259 filtered MSAs.

In *Brachypodium,* transcripts assembled *de novo* from RNA-seq data (supplementary tables S4 and S5), together with annotated transcripts or coding sequences (CDS) from *B. sylvaticum*, *B. distachyon*, *O. sativa* and *H. vulgare*, were clustered with GET_HOMOLOGUES-EST (Contreras-Moreira et al. 2017). Redundant and overlapping cluster sequences (at least 20 overlapping nucleotides) were collapsed with the *annotate_cluster* tool, producing 3,675 multi-copy FASTA files. The starting MSAs consisted of 3,675 individual transcript alignments for five diploid [*B. distachyon* (Bdis)*, B. stacei* (Bsta)*, B. arbuscula* (Barb)*, B. pinnatum* (Bpin), *B. sylvaticum* (Bsyl)], seven polyploid [*B. mexicanum* (Bmex) 4x*, B. boissieri* (Bboi) 6x*, B. retusum* (Bret) 4x*, B. hybridum* (Bhyb) 4x*, B. rupestre* (Brup) 4x, *B. phoenicoides* (Bpho6 and Bpho422), 4x], and two diploid outgroup [*H. vulgare* (Hvul) and *O. sativa* (Osat)] samples. After running *trim_MSA_block* tool and trimAl, a total of 2,001 filtered MSAs were recovered.

***1.2.* *Building, rooting and sorting the phylogenetic trees***

Preliminary phylogenetic trees were constructed for each of the filtered MSAs with IQ-TREE (Nguyen et al. 2014). Trees were rooted with the most distant outgroup (*O. sativa*) and sorted in decreasing order of branch divergence with the *reroot_tree* tool. Unrooted trees (i.e. without *O. sativa*) were discarded. The analyses recovered 259 and 1,924 phylogenetic rooted and sorted trees for the *Triticum*-*Aegilops* and *Brachypodium* groups, respectively.

***1.3. Selecting congruent topologies of diploid species in each phylogenetic tree***

Topologies containing only diploid species were inspected in pruned trees, keeping diploid ingroup and outgroup species and using the *check_diploids* tool that estimated the position of each diploid branch in each tree. The most frequent diploid topologies were *(O. sativa,(B. distachyon,(H. vulgare,((A. speltoides,(A. sharonensis, A. tauschii)),(T. monococcum, T. urartu)))))* for the *Triticum*-*Aegilops* group, and *(O. sativa,(H. vulgare,(B. stacei,(B. distachyon,(B. arbuscula,(B. pinnatum, B. sylvaticum))))))* for the *Brachypodium* group. These predominant diploid skeleton trees were recovered in 48 out of 259 (18%) filtered *Triticum*-*Aegilops* MSAs and 329 out of 1,924 (17%) *Brachypodium* MSAs. All the MSAs that showed the predominant diploid skeleton topology (including all their respective swapping-sister branch variants) were saved and used in downstream analyses.

We validated the selection of MSAs showing the most frequent diploid skeleton trees using coalescence-based ASTRAL, STAR and STEAC approaches that accounted for the potential existence of incomplete lineage sorting (ILS). All the coalescent analyses recovered the same optimal diploid species tree for, respectively, the *Triticum*-*Aegilops* (supplementary fig. S2A, B, C) and *Brachypodium* groups (supplementary fig. S4A, B, C).

***1.4. Labelling homeologous sequences (homeologs) of polyploid species***

Polyploid homeolog graftings in the selected MSAs’ trees that showed the congruent diploid skeleton topology were scrutinized to label these homeologs using the *check_lineage_polyploid* tool that labels them according to their grafting position with respect to ancestor, descendant and sister diploid lineage branches. For it, *ad hoc* labelling rules were defined using the ‘a’-‘i’ lowercase letters to label each homeolog tip (figs. 2A and 3A).

In the 48 *Triticum*-*Aegilops* MSAs, label ‘a’ was assigned to the graftings in the *Triticum*-*Aegilops* stem branch, ‘b’ to the *Aegilops*’s clade stem branch, ‘c’ to the *A. speltoides* branch (sister lineage), ‘d’ to the *A. sharonensis/A. tauschii* clade stem branch, ‘e’ to the *A. sharonensis* branch, ‘f’ to the *A. tauschii* branch, ‘g’ to the *T. monococcum/T. urartu* clade stem branch, and ‘h’ and ‘i’ to the *T. monococcum* and *T. urartu* branches, respectively (fig. 2A). In the 329 *Brachypodium* MSAs, the label ‘a’ was assigned to the graftings in the *Brachypodium* stem branch, ‘b’ to the *B. stacei* branch, ‘c’ to the branch sister to the *B. stacei* branch, ‘d’ to the *B. distachyon* branch, ‘e’ to the branch sister to the *B. distachyon* branch, ‘f’ to the *B. arbuscula* branch, ‘g’ to the branch sister to the *B. arbuscula* branch, and ‘h’ and ‘i’ to the *B. sylvaticum* and *B. pinnatum* branches, respectively (fig. 3A).

The *check_lineage_polyploid* tool generated three output files for each group: (a) ‘Labelled MSAs 1’ (*FASTA* format), which included only complete homeolog gapped sequences; (b) ‘Labelled MSAs 2’ (*FASTA* format), which included complete homeolog gapped sequences and missing data (subgenomic sequences) of non-detected homeologs (‘a’ to ‘i’) (these files were eventually concatenated to build the large multi-gene MSA); and (c) ‘Labelled gene tree’, which included all the computed gene trees showing the labelled polyploid branches (*Newick* format). Preliminary statistical reports were generated with the *make_lineage_stats* tool to check the number of homeologs recovered in each polyploid species.

**2. ‘Bootstrapping Refinement’ algorithm**

The labelled homeolog sequences obtained with the previous algorithm were further tested by a second labelling round using bootstrapping threshold support values for each of the grafted homeologs. Inconsistently labelled homeologs were discarded and the resulting double-filtered MSAs and their statistical values were used in downstream analyses.

***2.1. Testing the support of the labelled homeologs through a bootstrapping approach***

All the alignments that included one homeolog plus all diploid sequences of each MSA at a time were analysed through 1,000 bootstrapping trees (boottrees) generated with IQTREE. One file with 1,000 bootstrapping trees was generated for each pruned MSA. The initial number of homeologs was 236 in the 48 *Triticum-Aegilops* MSAs and 1,965 in the 329 *Brachypodium* MSAs.

***2.2. Extraction, rooting and sorting of bootstrapping trees***

Every boottree was extracted from the appended tree file, rooted, and ladderized using the *bootstrap_label_stat* tool with the ‘rootonly’ option. This generated 236,000 (236 x 1,000) files of rooted bootstrapping trees in *Triticum*-*Aegilops* and 1,965,000 (1,965 x 1,000) in *Brachypodium*.

***2.3. Selecting congruent supported topologies of diploid plus one-homeolog samples from the bootstrapping trees***

Similarly to step 1.3, the diploid topologies of the generated boottrees were inspected using the *check_diploids* tool but this time for trees that included the diploid plus one grafted homeolog sequences. Only those trees congruent with the diploid skeleton topology were saved. Of these, the homeologs from pruned MSAs that did not reach a threshold of branch grafting in ≥10% of the generated bootstrapping trees were discarded (0 in *Triticum*-*Aegilops*, 177 in *Brachypodium*). Then, a set of 100 bootstrapping trees showing the congruent diploid skeleton topology and containing only filtered homeologs was randomly selected for each of the pruned and selected MSAs of *Triticum*-*Aegilops* (48 MSAs, 236 homeologs) and *Brachypodium* (323 MSAs, 1,788 homeologs) for downstream analyses.

***2.4. Concatenating MSAs of the one hundred bootstrapping trees with congruent diploid skeleton topology and re-labelling process***

MSAs of the selected 100 boottrees were saved into a single file per pruned MSA and the homeologs contained in them were re-labelled following the same protocol as in step 1.4 and using the *bootstrap_label_stats* tool to test for the accuracy of the previous labels. Incongruently labelled homeologs (13 in *Triticum-Aegilops* and 383 in *Brachypodium*) in which the main bootstrapping label and the original label (step 1.4) did not match were discarded. A total of 223 polyploid homeologs from 48 MSAs and 1,405 from 322 MSAs remained, respectively, in the *Triticum*-*Aegilops* and *Brachypodium* data sets after this refinement.

***2.5. Removing underrepresented homeologs***

A threshold of ≤10% was applied to discard underrepresented homeolog graftings as they could be artefacts of the labelling process or non-informative subgenome-of-origin sequences (e. g. ancestral or highly variable residual variants). Statistical reports were generated using the *make_lineage_stats* tool. In the *Triticum*-*Aegilops* data set the highest percentages corresponded to the expected homeologs of allotetraploid *T. turgidum* (‘c’ 36.0%, ‘i’ 38.4%) and allohexaploid *T. aestivum* (‘c’ 25.5%, ‘f’ 32.8%, ‘i’ 27%) whereas the remaining homeologs’ graftings of each species showed percentages <10% (table 1A, homeolog-type). In the *Brachypodium* data set some allopolyploids showed different predominances of homeologs, like *B. mexicanum* [‘a’ (47.8%), ‘b’ (23.1%), ‘c’ (21.0%)], *B. boissieri* [‘a’ (39.5%), ‘b’ (17.5%), ‘c’ (22%), ‘e’ (12.4%)], and *B. retusum* [‘a’ (12.9%), ‘c’ (18.9%), ‘e’ (29.4%), ‘g’ (15.4%)] (table 1B, homeolog-type). The widely studied allotetraploid *B. hybridum*, which was used as a positive control, displayed similar percentages of homeologs ‘b’ (55.8%) and ‘d’ (42.5%), inherited from its respective diploid progenitors *B. stacei* and *B. distachyon* (table 1B, homeolog-type). The *Brachypodium* core perennial allotetraploids *B. rupestre* and *B. phoenicoides* (accessions Bpho6 and Bpho422) showed similar homeolog frequencies for the most recently evolved homeolog-types ‘e’ (21-25%), ‘f’ (12-20%), ‘g’ (23-29%), ‘h’ (12-20%) and ‘i’ (15-17%) (table 1B, homeolog-type). A total of 181 homeologs from 48 genes were retrieved for the *Triticum*-*Aegilops* data set and 1,307 homeologs from 322 genes for the *Brachypodium* data set (tables 1A, B).

***2.6. Homeologs’ ML consensus tree***

All labelled MSAs from step 2.5 (FASTA format) were concatenated with the *concat_alignment* tool from the GET_PHYLOMARKERS suite (Vinuesa et al. 2018), and consensus maximum likelihood (ML) phylogenetic trees of orthologs and selected homeologs were constructed with IQ-TREE for the respective *Triticum*-*Aegilops* (table 1A, Inferred Subgenome) and *Brachypodium* (table 1B, Inferred Subgenome) data sets. The annotated list of the 322 genes used to compute the *Brachypodium* homeologs’ ML consensus tree is shown in supplementary table S14.

**3. ‘Subgenome Assignment’ algorithm**

The *Subgenome Assignment* algorithm was used to infer the subgenomes of each polyploid. This algorithm selects the most frequent homeolog-types per polyploid according to its assigned subgenomic constitution inferred from the subgenomic definition steps and its ploidy. To define the subgenomes and assign the homeologs to them, we analyzed the grafting frequencies of the homeolog-types (‘a’-‘i’), computed across the 100 randomly selected bootstrapping relabelled trees of step 2.5. The homeologs’ grafting distributions were evaluated to determine their circumscription to a single or few contiguous branches of the species tree. The cytogenetic data (supplementary table S1) was used to inform the subgenomic assignments. In addition, we also employed a complementary phylogenetic criterion to improve the specificity of the subgenomic definition based on the topological resolution of the homeologs’ ML consensus tree of step 2.6 and the close proximity of homeolog-types in the first two axes of a Principal Coordinate Analysis (PCoA) and superimposed Minimum Spanning Tree (MST) generated from pairwise patristic distances from that tree.

***3.1a. Homeolog grafting distributions, subgenomic definition and assignment of homeologs to subgenomes***

To correct for the excess of homeologs in some polyploids and to assign them to defined subgenomes, we computed the relative frequencies (%) of homeologs detected in the studied polyploid species by our in-house *homeolog_distribution* tool across the selected MSAs and ranked the percentages of their grafting position (down-frequency ranks) across the 100 bootstrapping trees of step 2.4 (supplementary tables S2A, B and S6A, B).

The *homeolog_distribution* tool (R-script available at Github <https://github.com/eead-csic-compbio/allopolyploids>) is expressed as follows:

For all genes (**X=*1,2,..,n***) of a polyploid, for which a homeolog has been previously labelled according to its grafting position in a specific branch of the skeleton diploid tree (**J=a,b,c,d,e,f,g,h,i**) (see fig. 1) with our ‘*Nearest Diploid Species Node’* algorithm, given:

a) ***X_xj_***, the number of bootstrap replicates in which the homeolog of gene ***x*** was grafted into branch ***j***,

b)
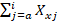
 = 100, the sum of bootstrap replicates in which the homeolog was grafted in all different **J** branches for gene ***x*** (100 in our study),

c) *S_j_* =
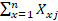
, the sum of bootstrap replicates in which the homeolog was grafted in branch ***j*** for all genes (**X**),

d) *S_tot_* =
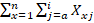
, the total number of graftings of the homeolog in all bootstrap replicates across all branches (**J)** and genes (**X**), and

e) *R****_j_*** *=* ***S_j_ / S_tot_***, the relative frequency of the total number of bootstrap replicates in which the homeolog was grafted in a specific branch ***j***,

the homeolog would be assigned to a simple or a compound genomic component (i. e., subgenome) based on the following criteria:

f) the homeolog grafted branch that defines a subgenomic group (***j = c***) is that showing the highest *R_j_* value *(****R_c_****)*. This *R****_c_*** homeolog grafted branch alone would define a simple subgenomic component (simple subgenome).

g) if other additional homeolog’s grafted branches ***( j ≠ c***) show ***R_j_ ≥ R_c_ / 10***, they would be included in a compound subgenomic component of the former (compound subgenome).

This algorithm consisted in the following steps: i) sorting homeologs’ grafting distributions downwardly by ranked frequency and circumscribing homeologs to single-type subgenomes if they were grafted to single-branches with the highest frequency and the remaining graftings were below a cut-off threshold (≤10% of the main grafting frequency), or to compound-type (multi-branch) subgenomes if the secondary and subsequent grafting frequencies were above the cut-off threshold; ii) choosing the most frequent homeolog-types according to the expected number of subgenomes suggested by ploidy (supplementary table S1; Marcussen et al. (2014a)); iii) re-coding the most frequent homeolog-types (‘a’, ‘b’, …) as subgenomes with capital letters (‘A’, ‘B’, …); iv) discarding low frequency homeolog-types incompatible with the ploidy level of the polyploid (if two or more most frequent homeolog-types were assigned to alternative single or compound subgenomes, then the less frequent homeolog-types were discarded and the process returned to step (i) with the remaining homeolog-types).

***3.1b. Homeologs’ ML consensus tree, Pairwise patristic distances, Principal Coordinate Analysis - Minimum Spanning Tree, and assignment of homeologs to subgenomes***

Pairwise patristic distances among all the orthologs and homeolog-types of the *Triticum*-*Aegilops* and *Brachypodium* homeologs’ ML trees generated in step 2.6 were calculated with Geneious R11.1.5 ([https://www.geneious.com](https://www.geneious.com/)) (supplementary tables S3 and S7). Separate Principal Coordinate Analysis (PCoA) was performed using the patristic distance values of the *Triticum*-*Aegilops* and *Brachypodium* data matrices with NTSYS-pc v2.10j (Rohlf 2000), and minimum spanning trees (MST) were superimposed on each of the PCoA plots (supplementary figs. S3, S5). Homeolog-types were assigned to single or compound subgenomes according to their divergent or close phylogenetic positions in the ML trees (figs. 2B, 3B) and in the bidimensional PCoA+MST plots (supplementary figs. S3, S5). Close homeolog-types were clustered into compound subgenomes. The subgenomic assignments of the homeologs were compared with those obtained from the ranked grafting distributions that circumscribe them to a single or few contiguous branches of the species tree in step 3.1a.

In *Triticum*-*Aegilops* all the selected homeologs were assigned to single subgenomes that contained only one homeolog-type (‘c’ corresponded to subgenome C, present in *T. turgidum* and *T. aestivum*, ‘f’ to subgenome F present in *T. aestivum*, and ‘i’ to subgenome I present in both polyploids) (table 1A, Inferred Subgenomes; supplementary table S2; fig. S3).

In *Brachypodium* some polyploids showed an excess of homeolog-types that required to rank and select the most frequent types or to cluster some of them to retrieve their plausible subgenomes (table 1B, Inferred Subgenomes; supplementary table S6; fig. S5). The allotetraploid *B. hybridum*, contained homeologs ‘b’ and ‘d’ that corresponded to its respective single progenitor subgenomes B and D (our B subgenome is equivalent to subgenome S (*B. stacei*-type) and D equals subgenome D (*B. distachyon*-type) in the nomenclatural system of the *Brachypodium* genomes (Gordon et al. 2020)). The remaining *Brachypodium* polyploids showed an excess of homeolog-types. The grafting uncertainty created around the stem branch and first diverging branches of the *Brachypodium* homeologs’ ML consensus tree and the MST of its PCoA plot (fig. 3B; supplementary fig. S5) likely generated spurious excess of homeolog-types ‘a’ to ‘c’ for the *Brachypodium* polyploids with ancestral subgenomes (*B. mexicanum*, *B. boissieri*, *B. retusum* pro parte), suggesting that they could belong to an ancestral subgenome. Nonetheless, the clear divergence of the *B. mexicanum* ‘a’ homeolog-type from the *B. boissieri* and *B. retusum* ‘a’ homeolog-type in the homeologs’ ML consensus tree and the PCoA-MST plot (fig. 3B; supplementary fig. S5) supported their respective assignments to independent ancestral A1 and A2 subgenomes. The successive divergences of the intermediately evolved *B. retusum* ‘e’ homeolog-type and the *B. rupestre*/*B. phoenicoides* ‘e’ homeolog-type in the ML tree (fig. 3B) sustained their assignments to independent intermediate E1 and E2 subgenomes, respectively (table 1B, Inferred Subgenomes; supplementary table S6; fig. S5). The close and recently evolved core perennial clade ‘f’, ‘g’, ‘h’ and ‘i’ homeologs were assigned to a recent G subgenome (table 1B, Inferred Subgenomes; supplementary table S6; fig. S5).

***3.2. Subgenomic ML consensus trees***

Filtered homeolog-types from step 2.6 were collapsed for each polyploid according to the subgenome assignments and relabelled with their respective single or compound subgenomic labels resulting in the subgenomic consensus labelled MSA and used to construct the subgenomic ML consensus tree with IQ-TREE for the respective *Triticum*-*Aegilops* (fig. 2C) and *Brachypodium* (fig. 3C) data sets.

**Detailed material and methods**

**Sampling, genome size, chromosome counting and ploidy level estimations of *Brachypodium***

Individual plants from ten vouchered *Brachypodium* species and ecotypes [diploids *B. arbuscula*, *B. pinnatum* and *B. stacei*, and polyploids *B. boissieri, B. hybridum, B. mexicanum, B. phoenicoides* (Bpho6 and Bpho422 accessions), *B. retusum* and *B. rupestre*] collected in their native circum-Mediterranean, Eurasian and North American (Mexico) regions were studied. Additional data from vouchered diploids *B. distachyon* and *B. sylvaticum* was incorporated to the study. Our sampling covered all the recognized diploid species that represent the main evolutionary splits of the genus (Catalán et al. 2016; Díaz-Pérez et al. 2018) (supplementary table S1).

Leaves of vouchered adult plants growing in pots were used for genome size (GS) estimation through flow cytometry. Nuclear suspensions were prepared from 200 mg of leaf sample and leaf internal standard. The nuclear DNA content of *B. retusum* was calculated using nuclei isolated from young leaves of *Raphanus sativus* “Saxa” (1.11 pg/2C DNA; (Doležel et al. 2007)) and those of *B. arbuscula*, *B. boissieri*, *B. mexicanum*, *B. phoenicoides*, *B. pinnatum* and *B. rupestre* using *Lycopersicon esculentum* ‘Stupicke’ (1.96 pg/2C DNA; (Doležel et al. 2007)) as standards. Nuclei were stained with propidium iodide and samples were analyzed using a CyFlow Ploidy Analyser SYSMEX following the protocol of Doležel et al. (2007). At least 5,000 nuclei were analyzed per sample. Each sample (two replicates) was analyzed three times on different days. Only measurements with coefficient of variation < 3.5% were recorded. Chromosome counting was performed on meristematic cells of roots obtained from germinated seeds or from hydroponic culture following the protocol of Jenkins and Hasterok (2007). Chromosome staining was performed with the DAPI fluorescent marker (4’, 6-diamino-2 phenylindole) and counting was done using a Motic BA410 fluorescence microscope. Ploidy levels were inferred from chromosome counts (2n) and GS estimations performed in the same accessions used in our transcriptome study and through contrasted GS and 2n values obtained in conspecific accessions that showed similar values (supplementary table S1). GS and chromosome values of the three annual *Brachypodium* species correspond to those indicated in Catalán et al. (2012). Cytogenetic information retrieved from other authors is indicated in supplementary table S1.

**Experimental design, RNA isolation, library preparation and transcriptomic data of *Brachypodium***

In order to maximize the number of expressed transcripts we could sample, the individual plants were divided to produce four genetically identical plants and each plant received four treatments [control (water every 48 h, 25ºC), soil drying stress (no water for one week), hot stress (40ºC day/25ºC night for 24 h), salt stress (500 mM NaCl in water, daily for two days)]. Total RNA was isolated from 50 – 200 mg of leaf tissue using the E.Z.N.A Plant RNA kit (Omega) or the RNeasy Plant Mini kit (Qiagen), following the manufacturers’ protocols. RNA integrity, purity and concentration were respectively measured with Agilent 2100 Bioanalyzer, NanoDrop, and Qubit, respectively. Pooled RNAs from the four treatments were used in subsequent RNA-Sequencing libraries. Library preparation was carried out by the Whitehead Genome Technology Core using TruSeq Stranded mRNA Library Prep Kit (Illumina, Inc.), generating Paired-End (PE) libraries with insert size of 300 to 600 bp. Library quantities were checked by Q-PCR and then sequenced using an Illumina HiSeq2500 platform at the Harvard University Bauer Core Facility.

Quality control of PE reads was performed with FastQC software (Andrews 2010). Adapters and low quality bases were removed and filtered with Trimmomatic-0.32 (Bolger et al. 2014). Transcript sequences were assembled with trinityrnaseq-r20140717 (Grabherr et al. 2011) using default parameters. *De novo* assembly of *Brachypodium* RNA-seq reads produced 72 to 160 thousand transcript isoforms with median lengths ranging between 414 to 555 bp (supplementary tables S4-S5).

*Brachypodium* RNA-seq data were deposited at ENA (European Nucleotide Archive; https://www.ebi.ac.uk/ena) with run accessions numbers ERR3633153 (*B. arbuscula*), ERR3634426 (*B. boissieri*), ERR3634464 (*B. hybridum*), ERR3634593 (*B. mexicanum*), ERR3634717 (*B. phoenicoides* Bpho422), ERR3634894 (*B. phoenicoides* Bpho6), ERR3634970 (*B. pinnatum*), ERR3636695 (*B. retusum*), ERR3636791 (*B. rupestre*) and ERR3636828 (*B. stacei*). *B. sylvaticum* RNA-seq data of the accession Brasy-Esp were obtained from the study by (Fox et al. 2013). Transcriptomic data was also retrieved for the relatively close *Oryza sativa* (SRX738077) and *Hordeum vulgare* (ERR159679) outgroup species that were used to root the phylogenetic *Brachypodium* trees and to provide a stem branch for the grafting of ancestral homeologs/subgenomes.

**Genomic data of *Triticum-Aegilops***

Genomic sequence data of *Triticum* and *Aegilops* species with known chromosome numbers and ploidy levels were retrieved from Marcussen et al. (2014a) and Marcussen et al. (2014b). It included 275 orthogroups of diploids *T. urartu*, *T. monococcum*, *A. speltoides*, *A. tauschii* and *A. sharonesis*, and allohexaploid *T. aestivum*. Additionally, cDNA sequences of the allotetraploid *T. turgidum* (*T. turgidum* subsp. *durum*; Svevo.v1; release 47 of <https://plants.ensembl.org/Triticum_turgidum/Info/Index>; (Maccaferri et al. 2019)) and genome data of the close outgroups *O. sativa* (Osativa_323_v7.0; http://phytozome.jgi.doe.gov/; (Ouyang et al. 2007)), *B. distachyon* (Bdistachyon_314_v3.1; <http://phytozome.jgi.doe.gov/>; (IBI 2010)) and *H. vulgare* (ftp://ftpmips.helmholtz-muenchen.de/plants/barley/genome_release2017/; (Mascher et al. 2017)) were added to this data set.

**Phylogenomic and dating analyses of the *Brachypodium* and *Triticum-Aegilops* data sets**

The alignments of the *Brachypodium* and *Triticum*-*Aegilops* data sets were conducted using GET_HOMOLOGUES-EST v09112017 (Contreras-Moreira et al. 2017) and MAFFT v7.222 (Katoh et al. 2002; Katoh and Standley 2013), respectively. Maximum Likelihood analyses were performed with IQ-TREE v.1.6.1 (Minh et al. 2013; Nguyen et al. 2014; Chernomor et al. 2016; Kalyaanamoorthy et al. 2017). The best-fit evolutionary model of each data set was automatically selected by ModelFinder (Kalyaanamoorthy et al. 2017) using the Akaike Information Criterion corrected (AICc). Topological congruence among alternative tree pairs was tested through Likelihood Ratio Test (SH-aLRT), and branch support ultrafast bootstrap searches were performed with 1000 replicates (Minh et al. 2013; Chernomor et al. 2016). The phylogenomic trees were conducted by “Partitioned analysis for multi-gene alignments” using the –spp option (Edge-proportional partition model with proportional branch lengths) of the IQ-TREE software.

In order to account for potential ILS in the selection of the most congruent diploid skeleton tree of each group, parallel analyses were conducted with the diploid sequences of the respective *Brachypodium* and *Triticum*-*Aegilops* data sets using ASTRAL v5.7.3 (Zhang et al. 2018), and STAR and STEAC (R v.3.5.1 package; (Liu and Yu 2010)) programs, which perform distance-based coalescence reconstructions.

Ancestral divergence ages of the *Brachypodium* homeologous subgenomes were estimated from a 322 nuclear core transcript data set (see Results) with BEAST 2.4.7 (Bouckaert et al. 2014). We imposed the GTR substitution model, lognormal relaxed clock (clock rate = 1x10^-4^) and Birth-Death tree models. We used two calibration points, imposing normal distribution secondary age constrains for the crown nodes of the BOP clade [*Brachypodium* + *Oryza* + *Hordeum*] (normal prior mean = 51.7 Ma, SD = 1.9) and the *Brachypodium* + core pooids clade [*Brachypodium* + *Hordeum*] (normal prior mean = 33.2 Ma, SD = 9.5), following the grass-wide plastome based dating analysis of Sancho et al. (2018), and a broad uniform distribution prior for the uncorrelated lognormal distribution (ucld) mean (lower = 1x10^-6^; upper = 0.1) and an exponential prior for ucld standard deviation (SD). The convergence of the parameters was reached at 582x10^6^ Markov chain Monte Carlo (MCMC) generations in BEAST2 with a sampling frequency of 5x10^3^ generations. The adequacy of parameters was checked using TRACER v.1.6 (<http://beast.bio.ed.ac.uk/Tracer>) with all the parameters showing Effective Sample Size (ESS) >200. Maximum clade credibility (MCC) trees were computed after discarding 10% of the respective saved trees as burn-in.

**Performance of the ‘Subgenome Assignment’ algorithm in the hypothetical presence of incomplete lineage sorting (ILS) in *Brachypodium***

Theoretical probabilities of each tested case (Strategy 1: simulated grafted allopolyploid subgenomes on tree branches; Strategy 2: simulated allopolyploids that matched true allopolyploids) were estimated by COAL across the 10,395 topologies that could be computed with 7 tips. The performance of the subgenome assignment algorithm was tested through all ranges of probabilities in each of the two strategies. In Strategy 1 the highest probability corresponded always to the species tree branch in which the homeologous subgenome was grafted to by our *Subgenome Assignment* algorithm. In Strategy 2 the *Subgenome Assignment* algorithm recovered the expected placements of the mimic allopolyploid subgenomes, despite the different topological graftings. In our Strategy 1 we also tested the alternative distributions of *B. retusum* [A2 (c)+E1 (e)] and *B. rupestre* [E2 (e)+G (h)] to corroborate if contiguous topological graftings of a subgenome could affect the effectiveness of the algorithm (supplementary table S9). Despite some homeolog grafting alternatives could be increased by ILS, as shown by these alternative tested distributions, they were discarded by the algorithm, indicating that grafting redundancies of the same subgenome in other close branches could be considered as artefacts.
